# Context-based facilitation in visual word recognition: Evidence for visual and lexical but not pre-lexical contributions

**DOI:** 10.1101/410795

**Authors:** Susanne Eisenhauer, Christian J. Fiebach, Benjamin Gagl

## Abstract

Word familiarity and predictive context facilitate visual word processing, leading to faster recognition times and reduced neuronal responses. Previously, models with and without top-down connections, including lexical-semantic, pre-lexical (e.g., orthographic/ phonological), and visual processing levels were successful in accounting for these facilitation effects. Here we systematically assessed context-based facilitation with a repetition priming task and explicitly dissociated pre-lexical and lexical processing levels using a pseudoword familiarization procedure. Experiment 1 investigated the temporal dynamics of neuronal facilitation effects with magnetoencephalography (MEG; N=38 human participants) while Experiment 2 assessed behavioral facilitation effects (N=24 human participants). Across all stimulus conditions, MEG demonstrated context-based facilitation across multiple time windows starting at 100 ms, in occipital brain areas. This finding indicates context based-facilitation at an early visual processing level. In both experiments, we furthermore found an interaction of context and lexical familiarity, such that stimuli with associated meaning showed the strongest context-dependent facilitation in brain activation and behavior. Using MEG, this facilitation effect could be localized to the left anterior temporal lobe at around 400 ms, indicating within-level (i.e., exclusively lexical-semantic) facilitation but no top-down effects on earlier processing stages. Increased pre-lexical familiarity (in pseudowords familiarized utilizing training) did not enhance or reduce context effects significantly. We conclude that context based-facilitation is achieved within visual and lexical processing levels. Finally, by testing alternative hypotheses derived from mechanistic accounts of repetition suppression, we suggest that the facilitatory context effects found here are implemented using a predictive coding mechanism.

**Significance Statement:** The goal of reading is to derive meaning from script. This highly automatized process benefits from facilitation depending on word familiarity and text context. Facilitation might occur exclusively *within* each level of word processing (i.e., visual, pre-lexical, and/or lexical-semantic) but could alternatively also propagate in a *top-down* manner from higher to lower levels. To test the relevance of these two alternative accounts at each processing level, we combined a pseudoword learning approach controlling for letter string familiarity with repetition priming. We found enhanced context-based facilitation at the lexical-semantic but not pre-lexical processing stage, and no evidence of top-down facilitation from lexical-semantic to earlier word recognition processes. We also identified predictive coding as the most likely mechanism underlying within-level context-based facilitation.

## 1 Introduction

Efficient reading relies on automatized visual word recognition (Rayner, 1998), which in turn involves visual-perceptual, pre-lexical orthographic and phonological, and subsequent lexical-semantic processing levels (e.g., Carreiras et al., 2014; Coltheart et al., 2001). Efficiency in reading depends mainly on our familiarity with the units of language (e.g., Gagl et al., 2015; Zoccolotti et al., 2008) and on facilitation that arises from the predictive nature of linguistic contexts during natural reading (e.g., Kliegl et al., 2006). Contextual facilitation results in reduced brain activation, most prominently of the N400, a component of the event-related brain potential (ERP) peaking ∼400 ms after word onset. The N400 reduction has typically been interpreted as reflecting facilitated processing at the lexical-semantic level of linguistic representation (see Kutas and Federmeier, 2011; Lau et al., 2008; for reviews). In line with this assumption, computational models like the strictly bottom-up sequential model of Laszlo and Armstrong (2014) successfully account for context-dependent N400 reduction effects by allowing neuronal fatigue *within* processing levels. Alternatively, however, it has also been proposed that contextual information (e.g., at the lexical-semantic level) can facilitate earlier stages of word recognition in a recurrent, *top-down* manner (see, e.g., Carreiras et al., 2014, for review). Fig. 1a visualizes these competing accounts of context effects on word recognition. Thus, the current model architectures disagree on the implementation of context-based facilitation as either within a processing level or top-down from higher processing levels.

**Figure 1.**
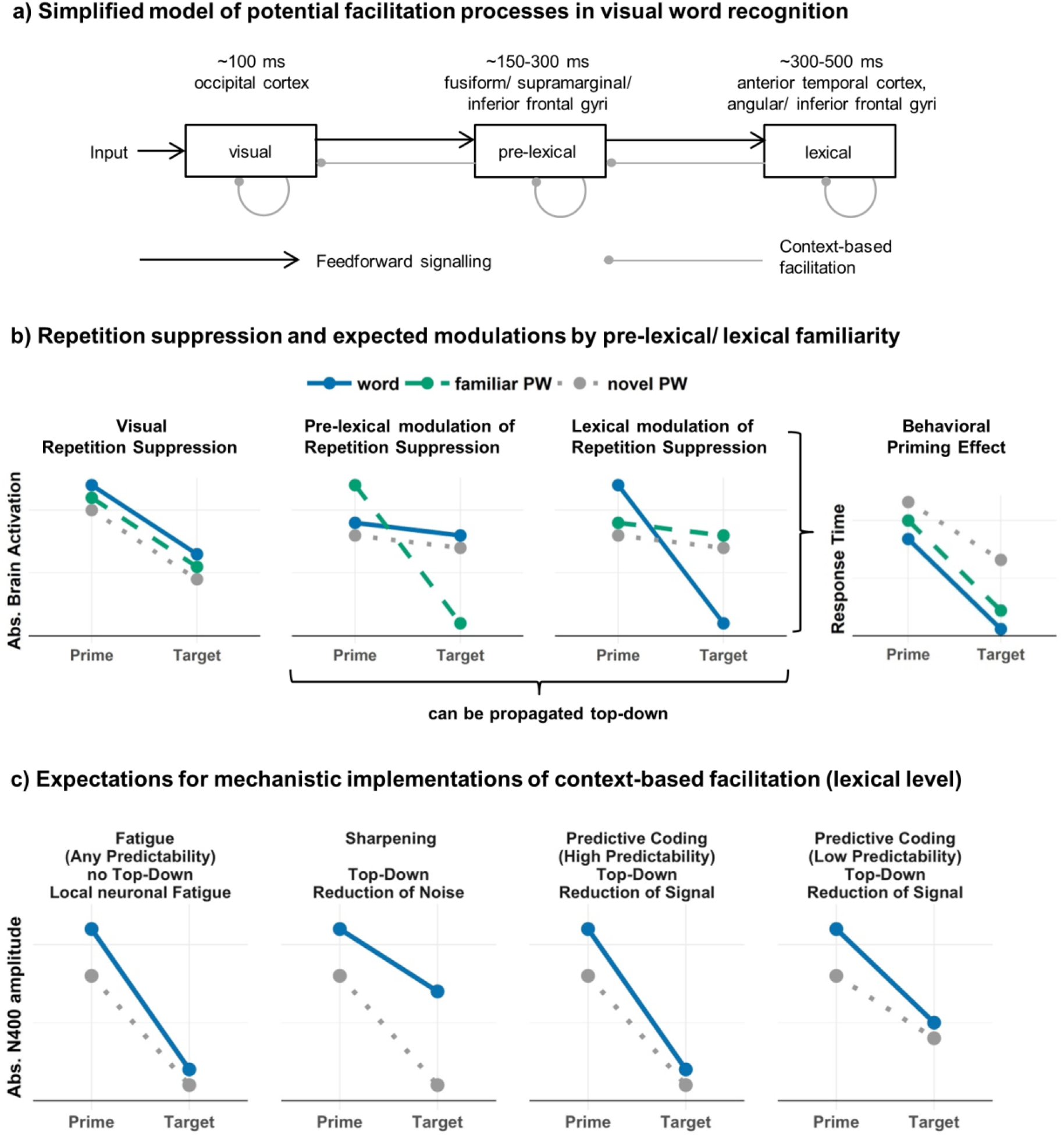
Model assumptions and predictions. a) A simplified architecture of visual word recognition (adapted from Carreiras et al., 2014) including expectations of ‘when’ (cf. Barber and Kutas, 2007) and ‘where’ in the brain the respective processes are implemented. Gray lines symbolize potential implementations of context-based facilitation either as within-level mechanism assumed in strictly bottom-up accounts (e.g., Laszlo and Armstrong, 2014) or, additionally, as a recursive top-down influence on hierarchically lower processing levels. b) Schematic representation of expected neuronal repetition suppression (left and central panels) and behavioral priming effects (right) with likely modulations by pre-lexical and lexical-semantic familiarity. We expect a reduction of neuronal activation and response times between primes and identical targets. At the lowest, i.e., visual processing level, we expect no familiarity modulation since all letter strings were a priori visually similar (left panel). For pseudowords (PW) that were familiarized through repeated exposure but without learning a new meaning, we expect selectively stronger neural repetition suppression in a ‘pre-lexical time window’ around 150 and 300 ms (central left panel). Finding this interaction additionally at earlier time points would be evidence for a facilitatory top-down influence of pre-lexical familiarity onto earlier visual processing stages. Note that as a manipulation check, we also expect that familiar pseudowords should elicit increased activation already at the prime at the expected regions (Gagl et al., 2016; Laszlo et al., 2014). At the level of lexical-semantic processing, we expect stronger repetition suppression for words compared to meaningless pseudowords (central right panel) at around 400 ms and, in case of top-down modulation, also in previous time windows. During prime processing, activation should be highest for words reflecting lexical-semantic processing (e.g., Rabovsky et al., 2012). In behavior, we expect stronger priming effects for words and familiar pseudowords compared to novel pseudowords, which would indicate that both pre-lexical and lexical-semantic familiarity increase context-based facilitation. c) Schematic visualization of expectations for fatigue, sharpening, and predictive coding mechanisms, shown for the N400 time window (i.e., lexical-semantic processing) and the contrast of words vs. novel pseudowords. The left panel shows an activation pattern reflecting a fatigue mechanism. This account assumes that the more activation is elicited on the prime (i.e., more semantic processing), the more neurons are ‘exhausted’, resulting in a stronger reduction for words vs. pseudowords. Sharpening (2^nd^ panel from left) expects a reduction of irrelevant (i.e., noisy) representations, thereby amplifying the signal. Consequently, neuronal repetition suppression should be weaker for words reflecting a focus of processing resources on informative word representations and, thus, should reduce activations selectively for pseudowords (e.g., Kok et al.; 2012). Predictive coding (two right panels) assumes a suppression of expected signals from the input. Thus, one would expect a suppression of the predictable signal rather than of the noise (e.g., Blank and Davis; 2016). For words, the additional lexical information can be used to predict the future target, resulting in stronger repetition suppression compared to pseudowords. The similar patterns for fatigue and predictive coding can be differentiated based on the probability with which identical prime-target pairs occur (see Grotheer and Kovács, 2014), which we implemented in Experiment 2. For predictive coding, one would expect stronger repetition suppression with higher repetition-probability, whereas the fatigue account makes no differential predictions depending on repetition-probability.

Computationally, Laszlo and Armstrong (2014) implemented context-based facilitation by a fatigue mechanism, assuming that recently active neurons are less likely to fire again (Grill-Spector et al., 2006). However, findings from semantic priming (e.g., Lau et al., 2013b) indicate an alternative mechanism, i.e., predictive coding, which assumes the suppression of perceptual signals that are consistent with context-based internally generated expectations about upcoming input (Friston, 2005). According to this model, one processes only the residual, i.e., unpredicted part of the input, which accounts for increased N400 activation when contextual expectations are violated. A third alternative would be sharpening (Grill-Spector et al., 2006), which assumes a reduction of neuronal firing only when neurons code the input suboptimally, thereby increasing the reliability of dissociating between inputs (e.g., Blank and Davis, 2016; Richter et al.; 2018). To date, we are not aware of a direct comparison of the possible mechanisms of context-based facilitation in visual word processing.

Separation of processing levels in visual word recognition research remains challenging. Experimental priming paradigms ensure a high degree of control over context factors, but require the matching of large numbers of psycho-linguistic parameters, which is difficult (Sassenhagen and Alday, 2016). Alternative regression-based accounts include these parameters as covariates, which can be realized in more natural contexts (i.e., sentence reading; e.g., Dambacher et al., 2006) but often demands large datasets (e.g., Dufau et al., 2015). We here propose that some of these problems can be ameliorated by using learning paradigms to increase familiarity and to associate information at different levels of linguistic representation with previously unfamiliar items (e.g., pseudowords: pronounceable non-words) in a controlled fashion (e.g., Taylor et al., 2011).

Using this strategy, we here dissociate between facilitation at pre-lexical and lexical levels of word processing. First, we matched pre-lexical characteristics (phonological length and orthographic familiarity) across words and pseudowords, ensuring comparable levels of pre-lexical processing difficulty between conditions (cf. Yarkoni et al., 2008). Second, we used a learning paradigm to increase pre-lexical familiarity of a subset of pseudowords (cf. Glezer et al., 2015) and measured highly time-resolved brain activation using magnetoencephalography (MEG; Experiment 1). In a second, behavioral, experiment, we also included pseudowords to which meaning was associated. To manipulate context-based facilitation, we used repetition priming (see DeLong et al., 2014, for a discussion of priming as context manipulation). Fig. 1b shows, in detail, the expected electrophysiological and behavioral responses reflecting context-based facilitation at visual, pre-lexical, and lexical-semantic processing levels. For example, we expect an interaction of lexical-semantic familiarity (presence vs. absence of lexical-semantic information) and context (prime/without context vs. target/with context) reflected by a stronger activation decrease (repetition suppression) for words in contrast to meaningless pseudowords (as shown by, e.g., Almeida and Poeppel, 2013). If restricted to the N400 time window, this pattern would indicate that facilitation is implemented exclusively at the lexical-semantic level, whereas earlier effects would suggest top-down facilitation from lexical-semantic to earlier processing stages. Source localization of these effects will help to specify these conclusions further. Finally, we aim at clarifying the mechanistic implementation of context-based facilitation, by comparing predictions from predictive coding, sharpening, and fatigue (see Fig. 1c).

## 2 Experiment 1: Magnetoencephalography

In the first experiment, we investigated pre-lexical vs. lexical-semantic contributions to context-based facilitation in visual word recognition at a neuronal level, using magnetoencephalography (MEG). Pre-lexical properties (orthographic Levenshtein distance/ OLD20; Yarkoni et al., 2008) were matched between words and both pseudoword groups (i.e., familiarized vs. novel), so that a priori, comparable levels of pre-lexical familiarity should lead to similar levels of pre-lexical activation across all three stimulus groups (as expected from, e.g., implementation of the MROM model: Grainger and Jacobs, 1996). However, the familiarization training increases the pre-lexical familiarity with the trained pseudowords. We expected effects of pre-lexical familiarity, i.e., increased activation of event-related fields (ERFs) for familiar in contrast to novel pseudowords (Gagl et al., 2016; Laszlo et al., 2014), and lexical familiarity, i.e., increased activation for words in contrast to novel pseudowords (cf. Rabovsky et al., 2012), irrespective of context. As an effect on context, we expected stronger neuronal repetition suppression for familiarized pseudowords compared to novel pseudowords and words on the pre-lexical level (see Fig. 1b central left panel) and stronger repetition suppression for words in contrast to the two pseudoword groups at the lexical level (see Fig. 1b central right panel). If we find this pattern, expected for pre-lexical processing, from 150-300 ms at, e.g., left fusiform regions, one can assume within-level context-based facilitation. One could come to a similar conclusion for lexical processing when the interaction pattern described above is found within the N400 time window at, e.g., left anterior temporal regions. Finding a similar pattern at earlier time windows would indicate top-down context-based facilitation.

### 2.1 Methods

#### 2.1.1 Participants

38 healthy native speakers of German (26 females, mean age 23.0±2.8 years, range 18-29 years) recruited from university campuses participated in familiarization procedures and MEG recordings and were included in the final sensor level analyses. All participants were right-handed as assessed by the Edinburgh Handedness Inventory (Oldfield, 1971), had normal or corrected-to-normal vision, and normal reading abilities as assessed with an adult version of the Salzburg Reading Screening (unpublished adult version of Mayringer and Wimmer, 2003). 19 further participants were excluded at different stages of the experimental procedure, due to the following reasons: Low reading skills (i.e., reading test score below 16^th^ percentile; N = 5), insufficient performance during pseudoword familiarization (i.e., accuracy for to-be-familiarized pseudowords < 50 % in the final learning session; N = 2), self-reported developmental speech disorder (N = 1), technical artifacts during the MEG measurement (N = 4), insufficient number of trials after artifact rejection (i.e., participants with < 15 repetition trials in at least one condition, outliers defined as >1.5 times below the lower quartile range of the number of valid trials across all participants and familiarity conditions; N = 2), contraindication to MEG measurement (N = 1, participant with retainer which might cause artifacts in MEG data), or drop out by choice of participants (N = 4, participants did not finalize the experimental procedure). All participants gave written informed consent according to procedures approved by the local ethics committees (University Clinic of Goethe University Frankfurt, Germany; application N° 107/15 and Department of Psychology, Goethe University Frankfurt, Germany; application N° 2015-229) and received 10 € per hour or course credit as compensation.

#### 2.1.2 Stimuli and presentation procedure

Words and pseudowords consisted of five letters, with the first letter in uppercase following convention for German nouns. Pseudowords were generated by the *Wuggy* software (Keuleers and Brysbaert, 2010), conserving the phonological (i.e., sub-syllabic) structure of the input words; all pseudowords were pronounceable. Estimates of word frequency and orthographic Levenshtein distance 20 (OLD20; Yarkoni et al., 2008) were based on the SUBTLEX-DE database (Brysbaert et al., 2011). A complete list of stimuli including estimated variables will be made available at OSF.

Sixty German nouns (logarithmic word frequency: mean ± standard error = 2.14 ± 0.12, range 0.00 to 4.03) and 120 pronounceable pseudowords were presented twice during MEG acquisition. In addition, 80 catch trials were presented (see below). Pseudowords were divided into two sets of 60 items, such that both pseudoword lists and the set of words were matched on orthographic similarity (OLD20; group means: ± one standard deviation: 1.825 ± 0.013; 1.717 ± 0.026; 1.743 ± 0.027) and number of syllables (1.817 ± 0.050; 1.95 ± 0.028; 1.933 ± 0.032; see Table 1 for stimulus characteristics). Despite the high similarity of the word characteristics between groups, these characteristics were included in all post hoc linear mixed models (LMMs) to account for potential confounds from the parameters (see analysis section for details). Participants were familiarized with 60 pseudowords before the actual repetition priming task was conducted in the MEG (see details of the familiarization procedure below). The second group of pseudowords was never seen by the participants before the MEG experiment. In addition, four further lists of 120 pseudowords each were generated as fillers for the familiarization procedure (one list per session).

**Table 1.** OLD20 and number of syllables for conditions of Experiment 1 (words, familiar and novel pseudowords) and Experiment 2 (words and three pseudoword [PW] sets).

|  | Minimum | 1<sup>st</sup> Quartile | Median | 3<sup>rd</sup> Quartile | Maximum | Mean | SE |
| --- | --- | --- | --- | --- | --- | --- | --- |
| **Experiment 1: OLD20** |  |  |  |  |  |  |  |
| **Words** | 1.600 | 1.750 | 1.850 | 1.900 | 2.050 | 1.825 | 0.013 |
| **Familiar** | 1.250 | 1.600 | 1.725 | 1.863 | 2.100 | 1.717 | 0.026 |
| **Novel** | 1.250 | 1.637 | 1.750 | 1.863 | 2.300 | 1.743 | 0.027 |
| **Experiment 1: Number of syllables** |  |  |  |  |  |  |  |
| **Words** | 1.00 | 2.00 | 2.00 | 2.00 | 2.00 | 1.817 | 0.050 |
| **Familiar** | 1.00 | 2.00 | 2.00 | 2.00 | 2.00 | 1.95 | 0.028 |
| **Novel** | 1.00 | 2.00 | 2.00 | 2.00 | 2.00 | 1.933 | 0.032 |
| **Experiment 1: Logarithmic word frequency** |  |  |  |  |  |  |  |
| **Words** | 0.000 | 1.512 | 2.229 | 2.858 | 4.032 | 2.137 | 0.115 |
| **Experiment 2: OLD20** |  |  |  |  |  |  |  |
| **Words** | 1.000 | 1.288 | 1.650 | 1.750 | 1.950 | 1.538 | 0.038 |
| **PW set 1** | 1.000 | 1.500 | 1.650 | 1.762 | 2.000 | 1.605 | 0.032 |
| **PW set 2** | 1.000 | 1.288 | 1.650 | 1.850 | 2.100 | 1.542 | 0.045 |
| **PW set 3** | 1.000 | 1.337 | 1.700 | 1.850 | 2.300 | 1.596 | 0.044 |
| **Experiment 2: Number of syllables** |  |  |  |  |  |  |  |
| **Words** | 1.00 | 2.00 | 2.00 | 2.00 | 3.00 | 1.833 | 0.059 |
| **PW set 1** | 1.00 | 2.00 | 2.00 | 2.00 | 2.00 | 1.95 | 0.028 |
| **PW set 2** | 1.00 | 2.00 | 2.00 | 2.00 | 2.00 | 1.967 | 0.023 |
| **PW set 3** | 1.00 | 2.00 | 2.00 | 2.00 | 2.00 | 1.9 | 0.039 |
| **Experiment 2: Logarithmic word frequency** |  |  |  |  |  |  |  |
| **Words** | 0.000 | 1.518 | 1.971 | 2.189 | 3.301 | 1.933 | 0.095 |

*Note.* SE = standard error of the mean.

Stimuli were presented using Experiment Builder software (SR-Research Ltd., Ottawa, Ontario, Canada). Words and pseudowords were presented in black bold Courier New font (14 pt.) in front of a white background. In the behavioral sessions, stimuli were presented on an LCD monitor with a refresh rate of 60 Hz, while during the MEG session, stimuli were projected with a refresh rate of 60 Hz onto a translucent screen.

#### 2.1.3 Pseudoword Familiarization

Participants visited the lab on the two days prior to the MEG experiment, and during each visit completed two familiarization sessions of about 20 min length. The two previous days were chosen to take advantage of sleep consolidation effects (James et al., 2017). Each familiarization session started with reading aloud the pseudowords from a printed list. Reading errors were documented (mean across all sessions: 0.7 %). Subsequently, participants performed a computer-based old/new recognition task in which the to-be-familiarized pseudowords were presented two times per session, randomly intermingled with a new set of 120 filler pseudowords for every session (total of 480 filler pseudowords across all four sessions). For every pseudoword, participants had to indicate by button press as fast and accurately as possible, if it was familiar to them or not. Pseudowords were preceded by two black vertical bars displayed above and below the center of the screen where participants were asked to fixate (500 ms; Fig. 2a), and presentation was terminated with the button press.

Linear mixed model (LMM) analyses with session (centered and z-transformed) as fixed effect and participant and item as random effects on the intercept were performed with the *lme4* package (Bates et al., 2015) in R, version 3.4.1, 2017-06-30 (R Development Core Team, 2008). All effects with t > 2, reflecting that the effect differs from zero by more than two standard errors, were considered significant (note that p-values cannot be computed in a reasonable way in the LMM approach; see, e.g., Kliegl et al., 2011). Note that for one participant, data of Session 3 and 4 could not be saved due to technical issues. Old/new response sensitivity indices d’ (Green and Swets, 1966) significantly increased across familiarization sessions from 1.15 in Session 1 to 2.96 in Session 4 (see Fig. 3a; *estimate* = 0.66, *SE* = 0.039, *t* = 17.17; pairwise tests between subsequent sessions: all *t’s* > 5; see Extended Data Fig. 3-1 for details). This demonstrated that participants improved across sessions in distinguishing between familiar and novel (i.e., filler) pseudowords. Participants reached a high performance in the final session, with accuracies ranging between 70.0 and 99.2 % for familiar and between 59.2 and 99.2 % for filler pseudowords (Fig. 3b). Based on the strong improvement in sensitivity and the high performance in the final session, we conclude that pre-lexical familiarization of the trained pseudowords was successful.

**Figure 2.**
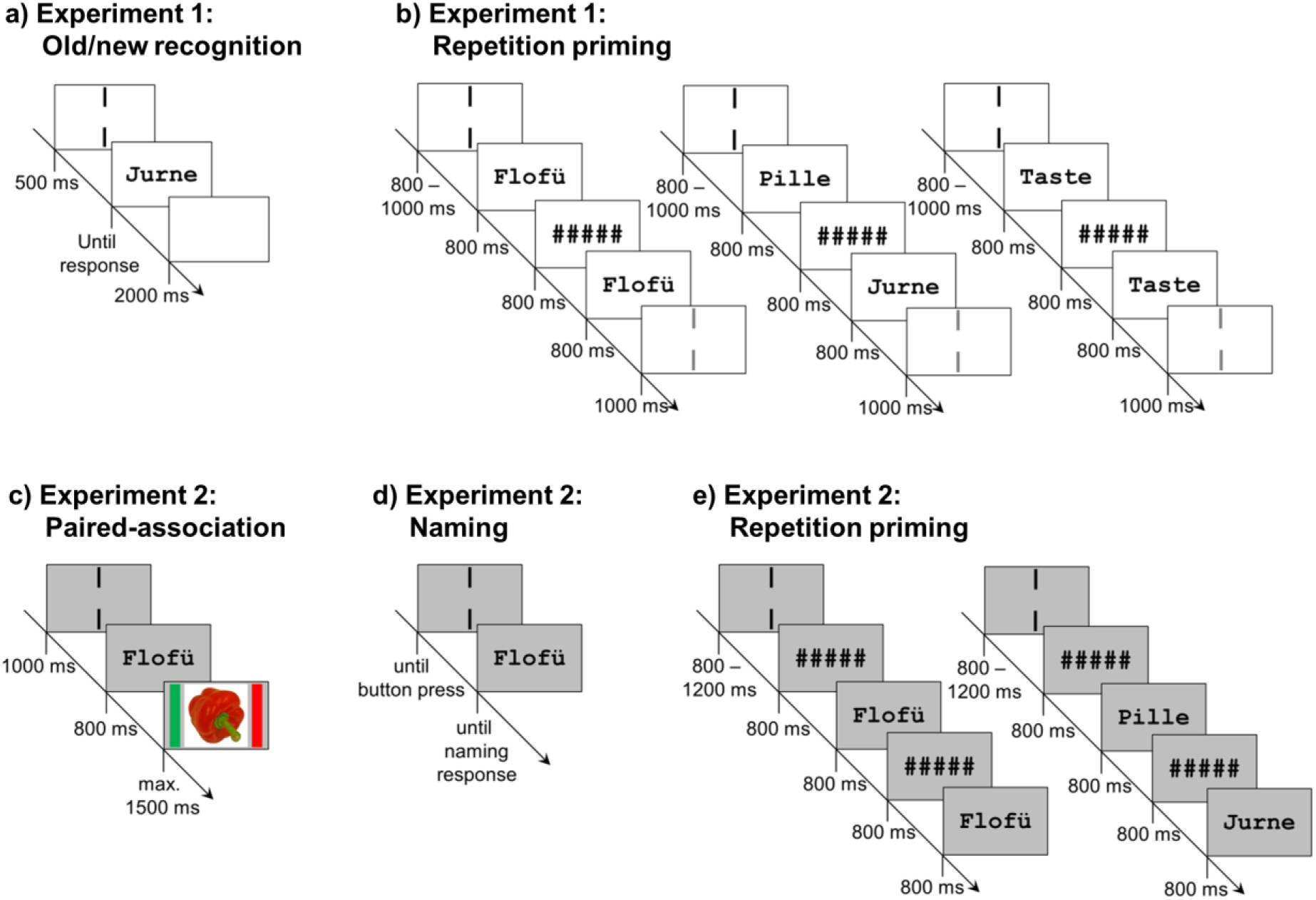
Experimental procedures. (a) For the pseudoword familiarization procedure of Experiment 1, in each learning session, 60 pseudowords were presented until response, intermingled by novel filler pseudowords, in an old/new recognition task. 500 ms before stimulus onset, two vertical bars indicated the center of the screen where participants were asked to fixate. The inter-trial interval was 2,000 ms. (b) During MEG recording, participants performed a repetition priming task. Each trial consisted of a sequence of two letter strings (prime and target) presented for 800 ms each, separated by an interval of 800 ms during which a string of five hash marks was presented. Letter strings could be words, familiarized pseudowords, or novel pseudowords (120 trials each regarding the prime). 75 % of trials were repetition trials, i.e. prime and target were identical (left). The remaining 25 % were non-repetition trials in which two different letter strings were presented (middle). In this case, prime and target could be from the same condition or from two different conditions, with all combinations of conditions appearing equally often. Participants were instructed to silently read presented letter strings and respond only to rare catch trials (right). Before onset of the prime, two black vertical bars presented for 800 – 1,000 ms indicated the center of the screen where participants were asked to fixate. After presentation of the target, two grey vertical bars were presented for 1,000 ms, indicating a blinking period of 1,500 ms starting from onset of the bars. Before the onset of the next trial, a blank screen was presented for the remainder of the blinking period. (c) In Experiment 2, a paired-association task was used for familiarization of pseudowords with and without semantics. Pseudowords were presented for 800 ms, followed by the presentation of an object image until button press (maximally 1,500 ms). During the inter-trial interval of 1,000 ms, two vertical bars indicated the center of the screen where participants were asked to fixate. In the semantic condition, there was a reliable association between object and pseudoword. In the familiarization only condition, in contrast, pseudowords and objects were randomly paired so that each pair occurred only once. (d) In the subsequent naming task, each pseudoword from the familiarization conditions with and without semantic associations was presented once. Participants named the object they associated with each pseudoword, or responded “next” in case they did not associate a meaning with a pseudoword. Before each pseudoword presentation, two vertical bars framing the center of the screen were presented until button press by the experimenter. (e) The repetition priming task involved in each trial a sequence of two letter strings presented for 800 ms each, separated by an interval of 800 ms during which five hash marks were displayed. The hash mark string was also presented for 800 ms before the onset of the first letter string. Letter strings could be words, familiarized pseudowords with and familiarized pseudowords without semantics, or novel pseudowords (180 trials each regarding the prime). Repetition probability was varied across blocks between 25, 50, and 75 %. Participants were instructed to silently read presented letter strings and respond to the target whether they had an explicit semantic association with it, or not. During the inter-trial interval of 800 – 1,200 ms, two vertical bars indicated the center of the screen where participants were asked to fixate.

**Figure 3.**
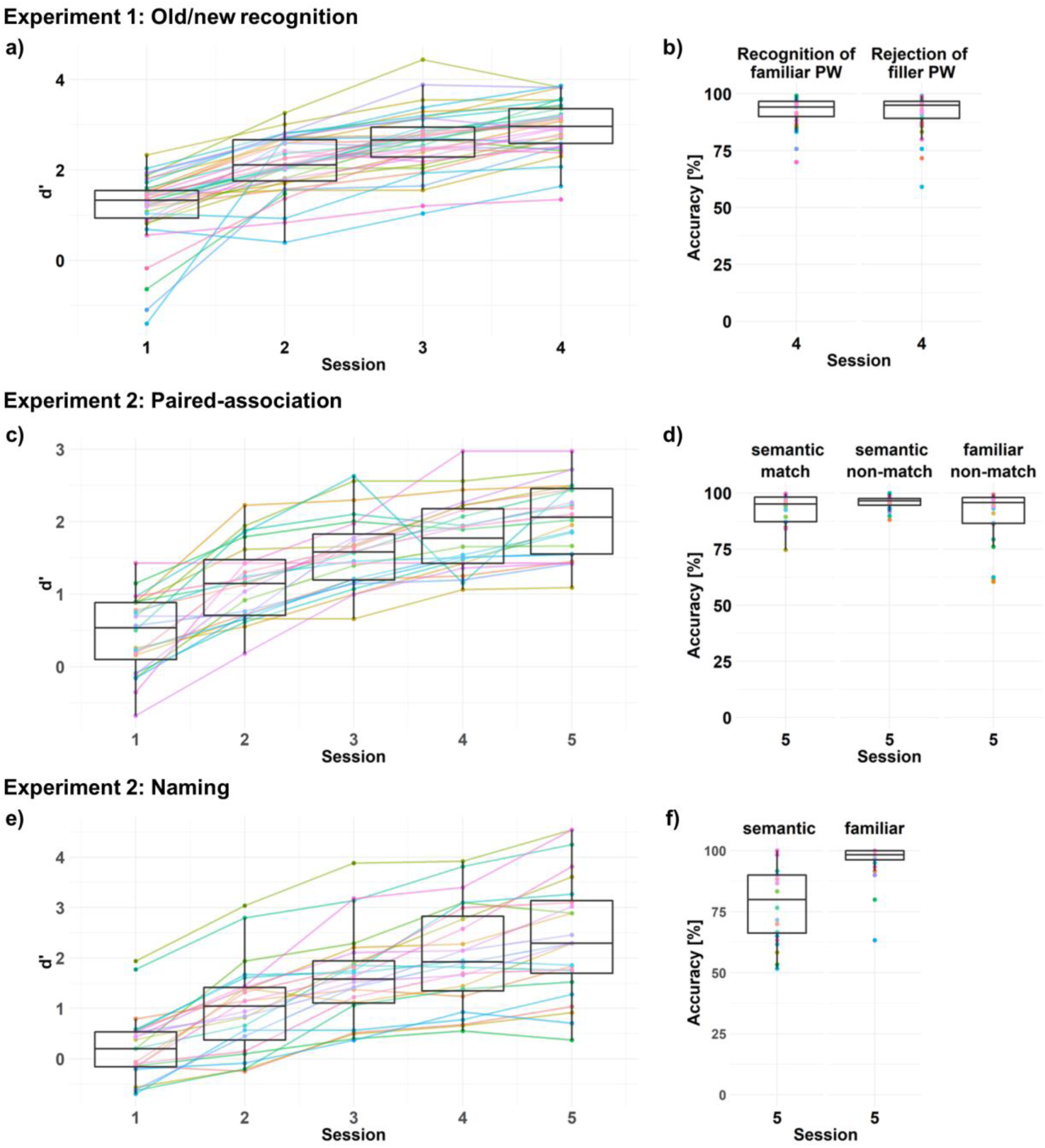
Behavioral results of familiarization procedures. a), b) Old/new recognition task (Experiment 1). c), d) Paired-association task (Experiment 2). e, f) Naming task (Experiment 2). *Left*, sensitivity indices d’ across all sessions. *Right*, accuracy in the final session. Colored dots and lines represent individual participants. Statistical results can be found in Extended Data Fig. 3-1.

#### 2.1.4 Repetition Priming

The repetition priming task during MEG recording was conducted on day 3 and included words, familiarized pseudowords, and novel pseudowords. At the start of each trial, participants had to fixate between two vertical black bars presented above and below the center of the screen (analogous to the familiarization procedure; cf. Fig. 2b). Stimulus presentation was initiated after an eye-fixation to the cued region was detected by an MEG compatible eye-tracker (Eyelink CL 1000, SR Research Ltd., Ottawa, ON, Canada), and comprised the successive presentation of two letter strings (prime and target) for 800 ms each, separated by an interval of 800 ms during which a string of five hash marks was shown. Both letter strings had to be read silently; the task served only to maintain attention and required a button press to catch trials, i.e., the word *Taste* (Engl.: *button*) in either the first, second, or both positions. The silent reading task was chosen to avoid contaminating the neuronal response to words with motor responses; catch trials were excluded from analysis. The explicit fixation control before stimulus presentation assured that eyes were open and directed towards the position where the stimulus was presented. Response hands were counterbalanced across participants and responses were recorded using a fiber optic response pad (LUMItouch; Photon Control Inc., Burnaby, BC, Canada). 100 ms after target offset, grey vertical bars were presented for 100 ms, indicating that participants were allowed to blink for a period of 1,000 ms. Stimuli were presented at a viewing distance of 51 cm yielding horizontal visual angles of about 0.3° per letter.

The 60 letter strings per condition (words, familiar, and novel pseudowords) were each presented in two trials, once during each half of the experiment. As a consequence, we presented 120 trials per stimulus condition adding up to 360 trials. 75% (i.e., 270) of these trials were repetition trials, allowing the investigation of familiarity effects in a highly predictive context. The remaining 25% (i.e., 90) trials were non-repetition trials, in which each possible combination of words, familiarized, and novel pseudowords appeared equally often, i.e., 10 times. Also, we presented 80 catch trials resulting in a total of 440 trials. The repetition priming task lasted about 40 min, divided into three blocks separated by breaks of about 5 min.

#### 2.1.5 MEG data acquisition

MEG datasets were acquired in accordance with guidelines for MEG recordings (Gross et al., 2013), using a 275 sensor whole-head system (Omega 2005; VSM MedTech Ltd., Coquitlan, BC, Canada). Six sensors (MLF66, MLP31, MRF22, MRF24, MRO21, and MZC02) were disabled due to technical issues so that 269 sensors remained for data acquisition. Data were recorded at a sampling frequency of 1,200 Hz using a synthetic third-order gradiometer configuration. Online filtering was performed with fourth-order Butterworth filters with 300 Hz low pass and 0.1 Hz high pass. Head positions of the participants relative to the gradiometer array were recorded continuously by three localization coils placed at the nasion and above both ear canal entrances using ear-plugs. Additionally, two electrodes placed centrally on each clavicula recorded an electrocardiogram (ECG), while two pairs of electrodes placed distal to the outer canthi of both eyes, and above and below the right eye, respectively, recorded an electrooculogram (EOG). The impedance of each electrode was below 5 kΩ for EOG electrodes and below 20 kΩ for ECG electrodes, measured with an electrode impedance meter (Astro-Med GmbH, Rodgau, Germany).

#### 2.1.6 Structural magnetic resonance image acquisition

Structural magnetic resonance (MR) images were acquired for 34 participants with a 1.5 T Siemens magnetom Allegra scanner (Siemens Medical Systems, Erlangen, Germany) using a standard T1 sequence (3D MPRAGE, 176 slices, 1 x 1 x 1 mm voxel size). To enable co-registration of MR images with MEG data, vitamin E capsules were placed at the positions of two of the MEG head localization coils (i.e., above both ear canal entrances using ear-plugs); the nasion could be identified anatomically in structural MR images. Fiducial coordinates were identified in SPM12 (http://www.fil.ion.ucl.ac.uk/spm/software/spm12/).

#### 2.1.7 MEG sensor level analyses

MEG data were analyzed with FieldTrip (Version 2011 11-21 for preprocessing and Version 2013 01-06 for all remaining sensor level analyses; http://fieldtrip.fcdonders.nl; Oostenveld et al., 2011) under MATLAB (version 2012b, The MathWorks Inc., Natick, MA), except for Fig. 4ab, which was realized with MNE-Python (https://martinos.org/mne/stable/index.html; Gramfort et al., 2013; 2014). Parallel computations were performed using GNU parallel (Tange, 2011). Catch trials and any other trials during which participants made a button press were excluded from analysis. MEG data were segmented into epochs of 2,600 ms length, lasting from −160 ms to 2,440 ms with respect to the onset of the prime.

**Figure 4.**
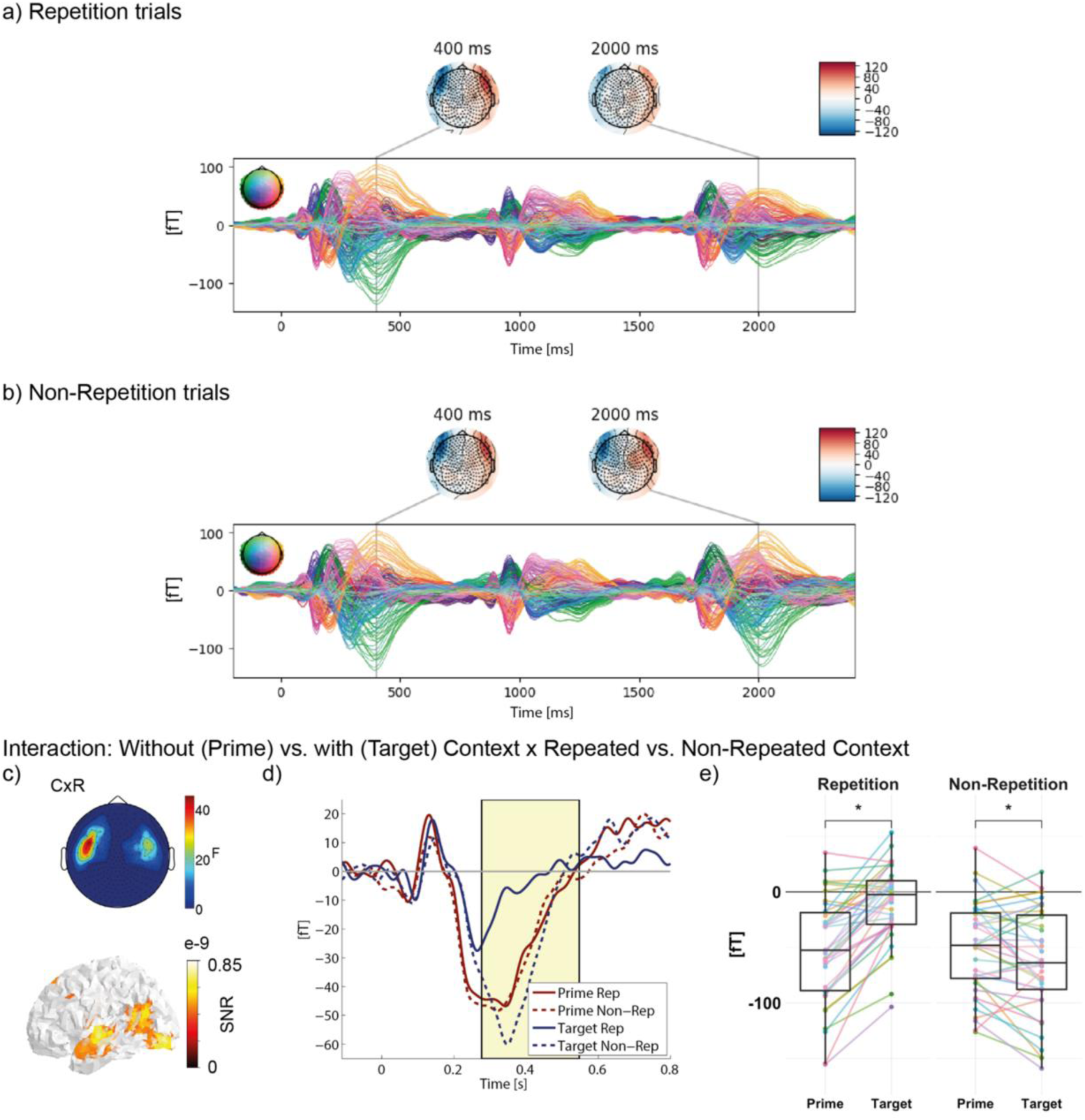
Repetition suppression: Interaction between trials without (prime) vs. with (target) context (C) and repeated vs. non-repeated context (i.e., repetition congruency; R). A detailed overview of all clusters obtained with common or separate baselines can be found in Extended Data Fig. 4-1. ERF time courses of a) repetition and b) non-repetition trials for all sensors and familiarity conditions. Colored lines represent individual sensors and within-plot topographical map color-codes scalp position of each sensor. Topographies represent activation at 400 ms and 2,000 ms, which allows a comparison of activation after 400 ms of the onset of prime and target, respectively. c) Topographical map represents *F*-values of significant sensors averaged across the significant time window. Non-significant sensors are set to zero. Surface plot represents source locations of the effect in signal-to-noise ratio (SNR) thresholded at 50 %. d) ERF time course averaged across significant sensors (of left hemisphere only, shown in topography in c). The significant time window is marked by a yellow shaded black box. Red lines correspond to prime and blue lines to target; solid lines correspond to repetition and dashed lines to non-repetition trials (averaged across all familiarity conditions). e) Boxplots represent activation averaged across sensors and time points within the left hemisphere cluster, for repetition trials (left) and non-repetition trials (right). Colored dots and lines represent individual participants. Asterisks indicate significant results (*t* > 2) from post hoc LMMs.

Individually for each participant, trials were selected for analysis in which the head position fell within a range of 5 mm (across all blocks) relative to the majority of other trials. Trials contaminated with sensor jump and muscle artifacts were rejected automatically, using the FieldTrip routine for automatic artifact detection. For jump artifact detection, a 9^th^ order median filter was applied to the data, while for muscle artifact detection, an 8^th^ order Butterworth IIR filter (110-140 Hz) was applied. The filtered data were z-transformed and averaged across sensors. Trials were rejected if for any time point the z value exceeded a threshold of z = 20 for jump artifacts and z = 6 for muscle artifacts, following standards established for the local measurement characteristics. Trials contaminated with eye blink, eye movement, or heartbeat artifacts were cleaned using Independent Component Analysis (ICA; Makeig et al.,1996). Components whose time courses correlated with EOG and ECG electrodes were rejected, using as threshold a correlation coefficient of r > 0.1, which sufficiently removed artifacts according to visual inspection. After these procedures, an average of 51.2 repetition trials (range 26 to 79) per condition could be retained. Non-repetition trials were averaged across conditions for analysis, with on average 52.6 trials per participant available (range: 29 to 80).

Prior to computation of ERFs, a 20 Hz low pass filter was applied to data epochs to increase the signal-to-noise ratio (cf. Lau et al., 2013a). In addition, to ensure that the low filter did not mask more transient components, we performed the main analyses after filtering at 40 Hz low pass. Original epochs were split into separate epochs for prime and target stimulus, ranging from −110 ms to 800 ms with respect to each stimulus onset. Epochs were baseline corrected by subtracting the average activation between −110 and −10 ms from each time point. For each sensor, we identified the participants for which the recorded magnetic field averaged across repetition trials and time lay outside the range of the mean across all participants ± 3.29 standard deviations; i.e. detecting extreme outliers outside the 99.9 % confidence interval. The signal of these noisy sensors (33 sensors in total; one to nine sensors in ten participants), per participant, was approximated by trial-wise interpolation from activation in neighboring sensors. ERFs were then calculated for each subject and condition (repeated words, repeated familiar pseudowords, and repeated novel pseudowords, as well as non-repetition trials combined across all familiarity conditions), separately for prime and target, by averaging the epochs across all trials. ERFs were compared between conditions using cluster-based permutation tests (Maris and Oostenveld, 2007) for dependent samples, corrected for multiple comparisons across time points (−110 to 800 ms) and sensors at cluster level. To compute interaction statistics, we used the permANOVA functions by Helbling (2015; https://github.com/sashel/permANOVA/). Clusters were defined as spatially and temporally adjacent samples with *F-*values exceeding an uncorrected α-level of 0.001 (cf. Eklund et al., 2016). The cluster-level statistic was calculated using the standard approach, i.e., taking the sum of *F-*values within a cluster (Maris and Oostenveld, 2007). Empirical cluster-level statistics were compared to the distribution of cluster-level statistics obtained from Monte Carlo simulations with 5,000 permutations, in which condition labels were randomly exchanged within each subject. Original cluster-level statistics larger than the 95^th^ percentile of the distribution of cluster-level statistics obtained in the permutation procedure were considered to be significant. Note, we use the terms ‘significant time window/ sensors’ for convenience. However, we are aware that the temporal and spatial extents of clusters obtained with the permutation procedure are subject to variations based on the signal to noise ratio, number of trials, and the selected cluster threshold. As a consequence, we will not interpret the exact values precisely and rather focus on the condition differences within the obtained clusters.

First, as a general check of our experimental manipulation, we assessed context effects by computing a 2×2 interaction between the experimental factors repetition congruency (repetition vs. non-repetition trials, reflecting whether context-based processing was indeed possible; referred to as R) and prime/target effect (reflecting the absence vs. presence of a preceding context; referred to as C). As the low number of non-repetition trials did not allow separate analyses of these effects for the different conditions, data were pooled across familiarity conditions (words, familiar, and novel pseudowords). Within each familiarity condition, the number of repetition trials was randomly stratified to match the number of non-repetition trials. In the second analysis, we examined in repetition trials how familiarity (words/lexical familiarity vs. familiar pseudowords/pre-lexical familiarity only vs. novel pseudowords) and context (prime vs. target) interact. This analysis served to examine the effects of different familiarity types on the neuronal repetition effect.

To determine the nature of significant interaction and main effects, we performed post hoc linear mixed model analyses for pairwise differences between relevant conditions. All post hoc tests were based on participant-and condition-specific ERF values averaged across sensors and time points from the respective significant cluster, and included participant and item as random effects on the intercept. Since not all trials entered the analyses due to exclusion of artefactual trials, which might have affected the matching across letter string conditions, OLD20 and number of syllables, both z-transformed and centered, were entered as additional fixed effects.

To rule out the possibility that our baseline correction approach, i.e., using separate baselines for ERFs elicited by prime and target stimulus in a trial, has created artificial effects due to the presentation of hash marks only before the target, we performed the analyses of repetition congruency by prime/target and familiarity by prime/target a second time, using the period before the prime as a common baseline for correction of ERFs to both stimuli. Of in total 24 significant clusters from the analyses after separate baselining, 17 were also found significant in the analysis after common baselining. Therefore, in the results and discussion sections, we will focus on those clusters replicated with the common baseline approach. Clusters with durations < 30 ms were not interpreted (cf. Dikker and Pylkkänen, 2013, for a similar approach), which led to the rejection of three clusters of ∼20 ms. A comparison of significant clusters from both analyses can be found in Extended Data Fig. 4-1 and 5-1.

As a further sanity check of the separate baselines approach, we additionally report a peak-to-peak analysis for the repetition by familiarity interactions as well as main effects. For this analysis, the positive (in case of right sensors) and negative (in case of left sensors) peaks of the ERFs were identified per participant, condition, and sensor (restricted to the time window +/-150 ms around the peak latency of grand average ERFs and the interval between 0 and 500 ms). In case of central sensors close to the midline (sensors MZC01, MZC03, MZC04, MZF01, MZF02, MZF03, MZO01, MZO02, MZO03, and MZP01), we separately decided whether to select the positive or negative peak, depending on which of the two peaks was absolutely higher in the across-participant ERFs. We decided against taking this approach in the majority of sensors because the ERFs typically declined during later time windows, in many cases reaching a value higher than the actual peak in absolute terms. Therefore, selecting the positive peak in the case of right sensors, and the negative peak in the case of left sensors, was the best compromise between automatic peak determination and avoidance of misplacing the actual peak value with a value that falls within the time range of decline of the ERF. We then subtracted the preceding peak value of respective other polarity (between stimulus onset and detected peak) from the already defined peak value. Statistical analyses were then performed on the absolute peak difference, using the cluster-based permutation procedure as described above, defining clusters solely based on spatial adjacency between sensors due to the lack of the temporal dimension. Given its independence from the pre-stimulus baseline, hash mark strings presented prior to the target cannot influence this analysis. However, a limitation of this analysis is that it cannot detect significant differences occurring at time ranges prior to and after the peak. Therefore, the results of this analysis did not influence whether a specific cluster was interpreted or not.

#### 2.1.8 MEG source localization

Source localization was performed for those 34 participants of whom we could obtain anatomical MR images, using FieldTrip, Version 2016 10-24. We created individual source grids for each participant by transforming the anatomical MR images to a standard (i.e., MNI space; Collins et al., 1994) T1 template from the SPM8 toolbox (http://www.fil.ion.ucl.ac.uk/spm). A regular 3D dipole grid (10 mm resolution) based on the T1 template was then warped with the inverse of the resulting transformation matrix. This procedure resulted in individual dipole grids for each participant, in which each grid point was located at the same brain area across participants. For each grid point and participant, lead fields were computed with a single shell forward model of the inner surface of the skull (Nolte, 2003). Prior to source localization, ERFs were down-sampled to 300 Hz to minimize computing costs. Source locations were computed for significant contrasts of interest from the ERF statistics as proposed by Gross et al. (2013). The procedure followed Manahova et al. (2018; the original code is available at https://data.donders.ru.nl/collections/di/dccn/DSC_3018012.15_439?0; see also Dijkstra et al., 2018; Mostert et al., 2018). Source localization was performed on ERFs by estimating two-dimensional dipole moments at each grid and time point using linearly constrained minimum variance (LCMV) spatial filters (van Veen et al., 1997). For the main prime/target effect, source localization was performed on the ERF difference between prime and target stimulus, averaged across all knowledge conditions in repetition trials. For main effects of familiarity, source localization was performed on the respective conditions separately and subtracted afterward (e.g., familiar-novel pseudowords averaged across prime and target). Interactions between repetition congruency and prime/target effect were resolved by performing source localization on the difference ERFs between stratified repetition and non-repetition trials, separately for prime and target, and then subtracting source activations of the prime from those of the target. Analogous, for interactions between prime/target and familiarity effects source localization was performed on the difference ERFs between prime and target, separately for each letter string condition, and then subtracted. The data covariance was estimated over the time interval of the respective significant contrast and regularized using shrinkage (Blankertz et al., 2011) with a regularization parameter of 0.01.

Two-dimensional dipole moments were reduced to a scalar value by taking the norm of the vector. This value reflects the contribution of a particular source location to sensor level activation not only in magnitude but also in dipole orientation. The latter is crucial for also detecting differences caused by different neuronal populations within the same source location. The norm of the vector is a positive value and subject to a positive bias due to noise. To counter this bias, we employed a permutation procedure (1,000 permutations). For analyses on separate conditions, the sign of half of the trials was randomly flipped. For analyses on condition differences, condition labels were randomly exchanged across trials. The square of the dipole’s norm averaged across all permutations was taken as noise estimate. This noise estimate was subtracted from the square of the true data, and the data were divided by the noise estimate. Negative values were set to zero, and the square root was taken. Finally, the signal of each source location was normalized by its variance in order to counter the depth bias. For visualization, source locations thresholded at 50 % of the maximum source activation were plotted on cortical surfaces using the *nilearn* package (Huntenburg et al., 2017) in Python. Brain regions were identified from the MNI coordinates of source maxima using the Harvard-Oxford cortical structural atlas (e.g., Desikan et al., 2006).

#### 2.1.9 Analysis code and data accessibility

The raw data, stimulus lists, and analysis code of both experiments described in the paper will be made freely available at OSF when published.

### 2.2 Results

During the MEG measurement, participants correctly identified 94 % of catch trials, indicating that they were attending to the presented letter strings.

#### Repetition suppression phenomenon

As manipulation check of context effects, we investigated the interaction between prime/target and repetition congruency (repetition vs. non-repetition trials) effects, combined over all familiarity conditions. Repetition trials (Fig. 4a) but not non-repetition trials (Fig. 4b) showed repetition suppression, i.e, reduced activity at the target stimulus (around 2,000 ms into the trial or 400 ms post onset of the target word). This interaction was significant at bilateral frontal sensors in the N400 time window (280 to 550 ms post-stimulus onset). Source localization revealed most prominently the left superior temporal gyrus (peak activation, extending into anterior temporal cortex albeit with weaker activation), as well as left occipital pole, left inferior occipital cortex, and the junction of left middle temporal, angular, and supramarginal gyri; Table 2; Fig. 4cd). The interaction reflected a significant decrease from prime to target in repetition trials (post hoc LMM: *estimate* = 2.22e^-14^, *SE* = 0.19e^-14^, *t* = 11.61; Fig. 4e left) and a significant but descriptively weaker increase in non-repetition trials (post hoc LMM: *estimate* = −0.58e^-14^, *SE* = 0.20e^-14^, *t* = 2.94; Fig. 4e right; see also Table 3). This replication of established repetition suppression effects (e.g., Deacon et al., 2004; Summerfield et al., 2011) is an important prerequisite for our main analyses investigating the effect of different familiarity conditions on repetition suppression. Sources of the effect were also compatible with previous localizations of the N400 within superior temporal gyrus (e.g., Helenius, 1998; Vartiainen et al., 2009); and anterior temporal cortex (e.g., Lau et al., 2013a; 2016; Lau and Nguyen, 2015).

**Table 2.**
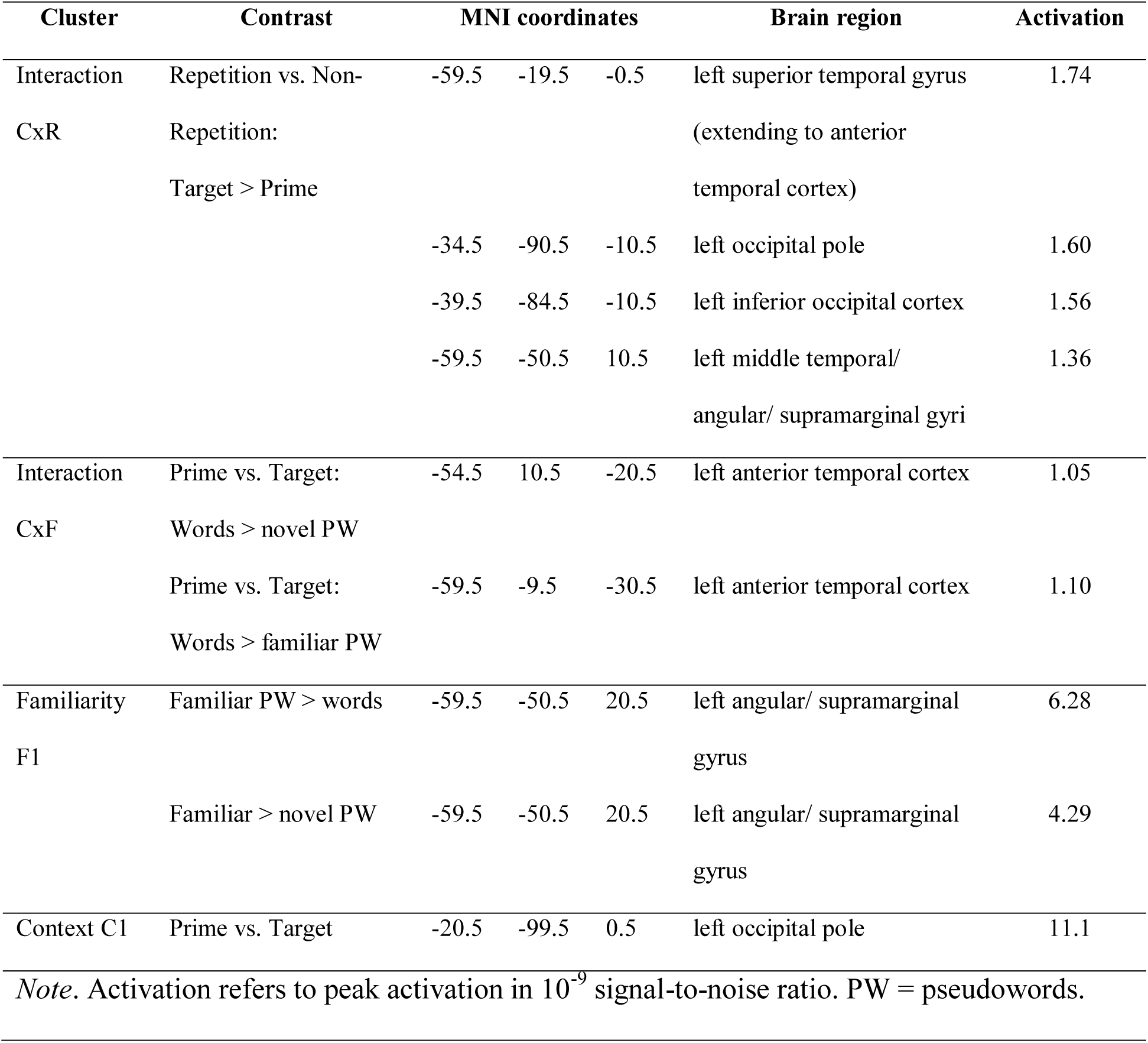
MNI coordinates of source maxima for clusters presented in Fig. 4, 5 and Extended Data Fig. 5-2.

**Table 3.** Results from post hoc linear mixed model analyses on ERF values (in 10^-13^ Tesla) from sensor and time point of the strongest effect for the prime/target x repetition congruency (repetition vs. non-repetition) interaction cluster represented in Fig. 4.

|  | Repetition |  |  | Non-Repetition |  |  |
| --- | --- | --- | --- | --- | --- | --- |
|  | *FE* | *SE* | *t* | *FE* | *SE* | *t* |
| Prime vs. target | **2.22** | **0.19** | **11.61** | **-0.58** | **0.20** | **2.94** |
| OLD20 | -0.18 | 0.22 | 0.81 | -0.28 | 0.25 | 1.09 |
| Number of syllables | 0.24 | 0.22 | 1.12 | -0.083 | 0.25 | 0.32 |

*Note.* Reliable effects are shown in bold. *FE* = fixed effects, *SE* = standard error.

#### Prime/Target effects

We here refer to context effects reflected in repetition suppression from prime to target in repetition trials only. Such effects were found in multiple time-windows, spanning time ranges from 100 to 690 ms after stimulus onset (Fig. 5a-d). All effects were localized to bilateral occipital cortices (see example of the earliest cluster C1 in Fig. 5a and source coordinates in Table 2). See also Fig. 5d for the sensor topographies of clusters C2-6. The majority of these clusters were not modulated by familiarity; only the frontal cluster C3 was qualified by a significant interaction with familiarity, indicated by the overlap in time and space with the interaction cluster (see below).

**Figure 5.**
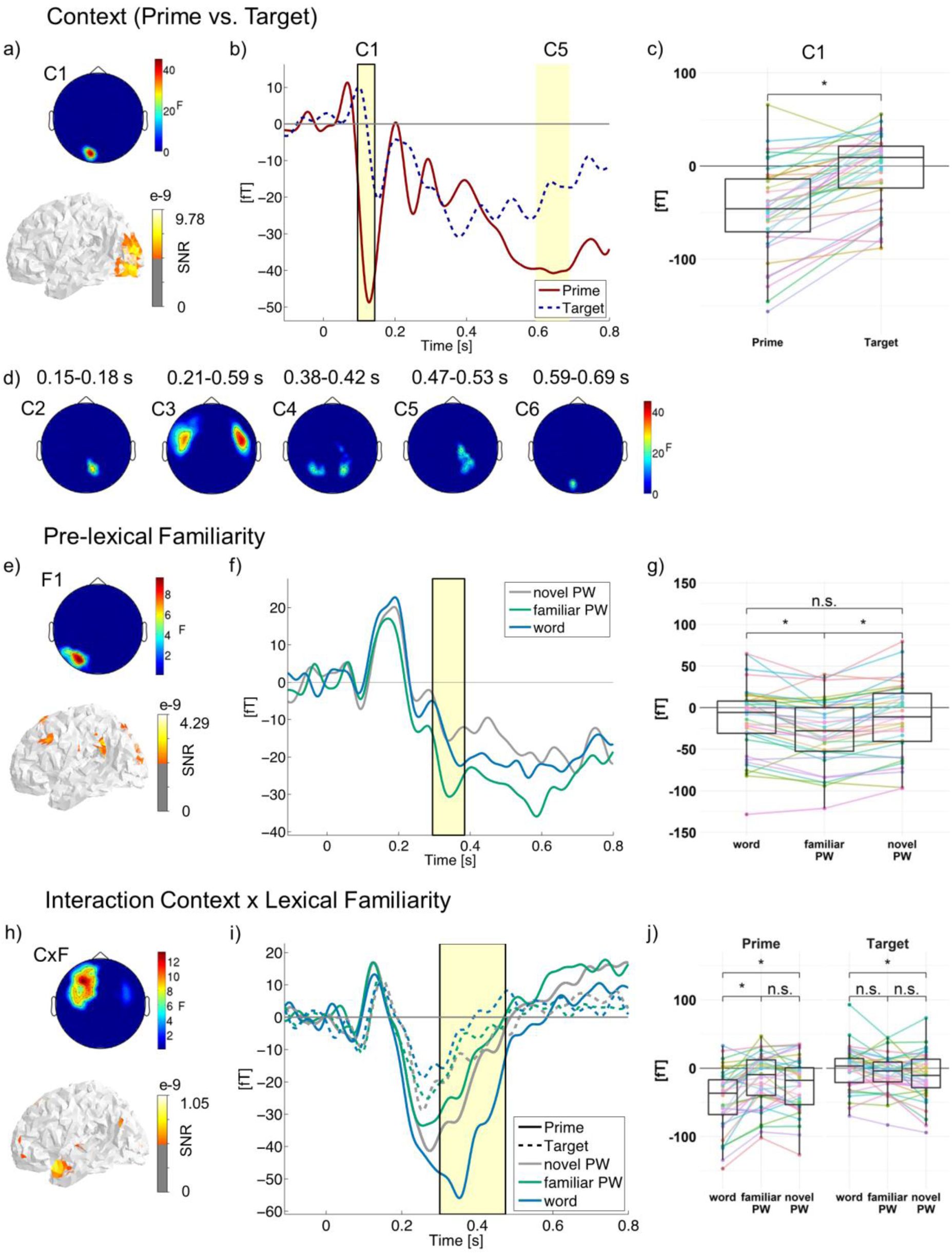
Main effects of context (prime vs. target; a-d), pre-lexical familiarity (e-g), and the influence of letter string familiarity on context effects (h-i; repetition trials only). A detailed overview of all clusters obtained with common baseline, separate baselines, and peak-to-peak analysis can be found in Extended Data Fig. 5-1. a) Topographical map represents *F*-values of sensors showing a main effect of context (prime vs. target) averaged across the significant time window. Non-significant sensors are set to zero. The surface plot represents the source location of the effect in signal-to-noise ratio (SNR) thresholded at 50 %. b) ERF time course averaged across the left occipital sensors (C1 which is marked by a black box; and C6 with similar sensor topography). The significant time windows are shaded in yellow. Solid red lines correspond to prime and dark blue dashed lines to target. c) Boxplots (from C1) represent activation averaged across sensors, time points, and familiarity conditions. Colored dots and lines represent individual participants. Asterisks indicate significant results (*t* > 2) from post hoc LMMs; n.s. = not significant. d) Additional topographical maps showing the main effects of context, which had similar effect patterns and source locations as C1. Significant time windows are indicated. e) Topographical map and source location (familiar > novel pseudowords, see Extended Data Fig. 5-2a for familiar pseudowords > words, f) ERF time course and g) boxplot showing a main effect of pre-lexical familiarity (i.e., differences between familiar pseudowords and words/ novel pseudowords averaged across prime and target). See Extended Data Fig. 5-3 for a main effect of lexical familiarity. h) Topographical map and source location (prime vs. target by words vs. novel pseudowords; see Extended Data Fig. 5-2b for prime vs. target by words vs. familiar pseudowords), i) ERF time course (left hemisphere only, shown in topography in h) and j) boxplots showing an interaction between context and lexical familiarity. Blue lines correspond to words, green lines to familiar and grey lines to novel pseudowords; solid lines correspond to prime and dashed lines to identical target. Note the different scales across contrasts. For results obtained with a low pass filter of 40 Hz instead of 20 Hz presented here, see Extended Data Fig. 5-4. Respective cluster topographies of the peak-to-peak analysis are shown in Fig. 5-5.

#### Familiarity effect

To reiterate, due to our stimulus selection procedure, words should have comparable levels of pre-lexical (orthographic/ phonological) familiarity but higher lexical familiarity (i.e., lexical-semantic information associated with words) than novel pseudowords. In contrast, as a result of the familiarization training, familiarized pseudowords should have higher levels of familiarity than novel pseudowords specifically at the level of pre-lexical processing. Thus, we had assumed that effects of pre-lexical familiarity should be reflected in ERF differences between familiar pseudowords and both novel pseudowords and words, while effects of lexical familiarity should be reflected in ERF differences between words and both novel and familiar pseudowords (cf. Fig. 1b for visualization of these hypotheses). Differences between the familiarity conditions, averaged across prime and target (i.e., representing main effects of familiarity), occurred at two topographic clusters: At left posterior sensors, localized to the left angular/ supramarginal gyrus (Fig. 5e, Extended Data 5-2a, Table 2), familiar pseudowords elicited a more negative ERF amplitude between 290 and 380 ms than both words and novel pseudowords (Fig. 5fg; Table 4, Extended Data Table 4-1). From the more negative ERF response specific to familiar pseudowords, we can conclude that our familiarization procedure was successful in modulating the neurophysiological processing of these pseudowords. The second cluster, again, occurred at frontal sensors and was qualified by a significant interaction with context (see below). The main effect in this cluster is therefore only visualized in Extended Data Fig. 5-3 (see also Table 4 and Extended Data Table 4-1) and not discussed further.

**Table 4.** Results from post hoc linear mixed model analyses on ERF values (in 10^-14^ Tesla) from sensor and time point of the strongest effect for prelexical (F1) and lexical (F2) familiarity clusters represented in Fig. 5e-g and Extended Data Fig. 5-4, respectively. Separate post hoc analyses for prime and target stimulus are represented in Extended Data Table 4-1.

|  | *F1* |  |  | *F2* |  |  |
| --- | --- | --- | --- | --- | --- | --- |
|  | *FE* | *SE* | *t* | *FE* | *SE* | *t* |
| Prime vs. target | **-0.59** | **0.12** | **4.71** | **2.87** | **0.17** | **17.33** |
| Pre-lexical familiarity | **-0.71** | **0.16** | **4.49** | **0.60** | **0.23** | **2.63** |
| Lexical familiarity | -0.080 | 0.16 | 0.49 | **-0.59** | **0.23** | **2.55** |
| Context x Pre-lexical familiarity | 0.079 | 0.14 | 0.55 | -0.20 | 0.19 | 1.04 |
| Context x Lexical familiarity | **-0.52** | **0.14** | **3.63** | **1.092** | **0.19** | **5.71** |
| OLD20 | -0.076 | 0.14 | 0.54 | -0.090 | 0.20 | 0.44 |
| Number of syllables | 0.0091 | 0.14 | 0.066 | -0.018 | 0.20 | 0.092 |

*Note.* Significant effects (i.e., $t > 2$ ) are shown in bold numerals. *FE* = fixed effect estimates, *SE* = standard error.

#### Pre-lexical and lexical familiarity effects on the repetition suppression phenomenon

We had expected that context effects reflected in repetition suppression for target relative to the identical prime be differentially influenced by pre-lexical vs. lexical familiarity (cf. Fig. 1b for visualization of these hypotheses). To assess the influence of familiarity on context effects, we examined the familiarity (words vs. familiar vs. novel pseudowords) by prime vs. target interaction in repetition trials (as only these included a valid predictive context for the target). We found a significant interaction between 300 and 480 ms at bilateral frontal sensors (Fig. 5hi; see also Table 5 for a post hoc statistic controlling for OLD20 and number of syllables). Post hoc LMMs revealed that lexical but not pre-lexical familiarity reliably modulated the repetition effect: While during prime presentation the negative-going ERF amplitude was largest for words (words vs. novel pseudowords: *estimate* = 2.15e^-14^, *SE* = 0.47e^-14^, *t* = 4.54; words vs. familiar pseudowords: *estimate* = 3.07e^-14^, *SE* = 0.42e^-14^, *t* = 7.39; no difference between familiar and novel pseudowords: *estimate* = −0.70e^-14^, *SE* = 0.40e^-14^, *t* = 1.75), it was smallest for words during target presentation (words vs. novel pseudowords: *estimate* = −1.20e^-14^, *SE* = 0.37e^-14^, *t* = 3.23; words vs. familiar pseudowords: *estimate* = −0.65e^-14^, *SE* = 0.37e^-14^, *t* = 1.76; familiar vs. novel pseudowords: *estimate* = −0.66e^-14^, *SE* = 0.33e^-14^, *t* = 1.98; see also Fig. 5j and Extended Data Table 5-1). Repetition suppression, thus, was stronger for words than for pseudowords. This effect was localized to the left anterior temporal cortex (Fig. 5h, Extended Data Fig. 5-2b, Table 2).

**Table 5.**
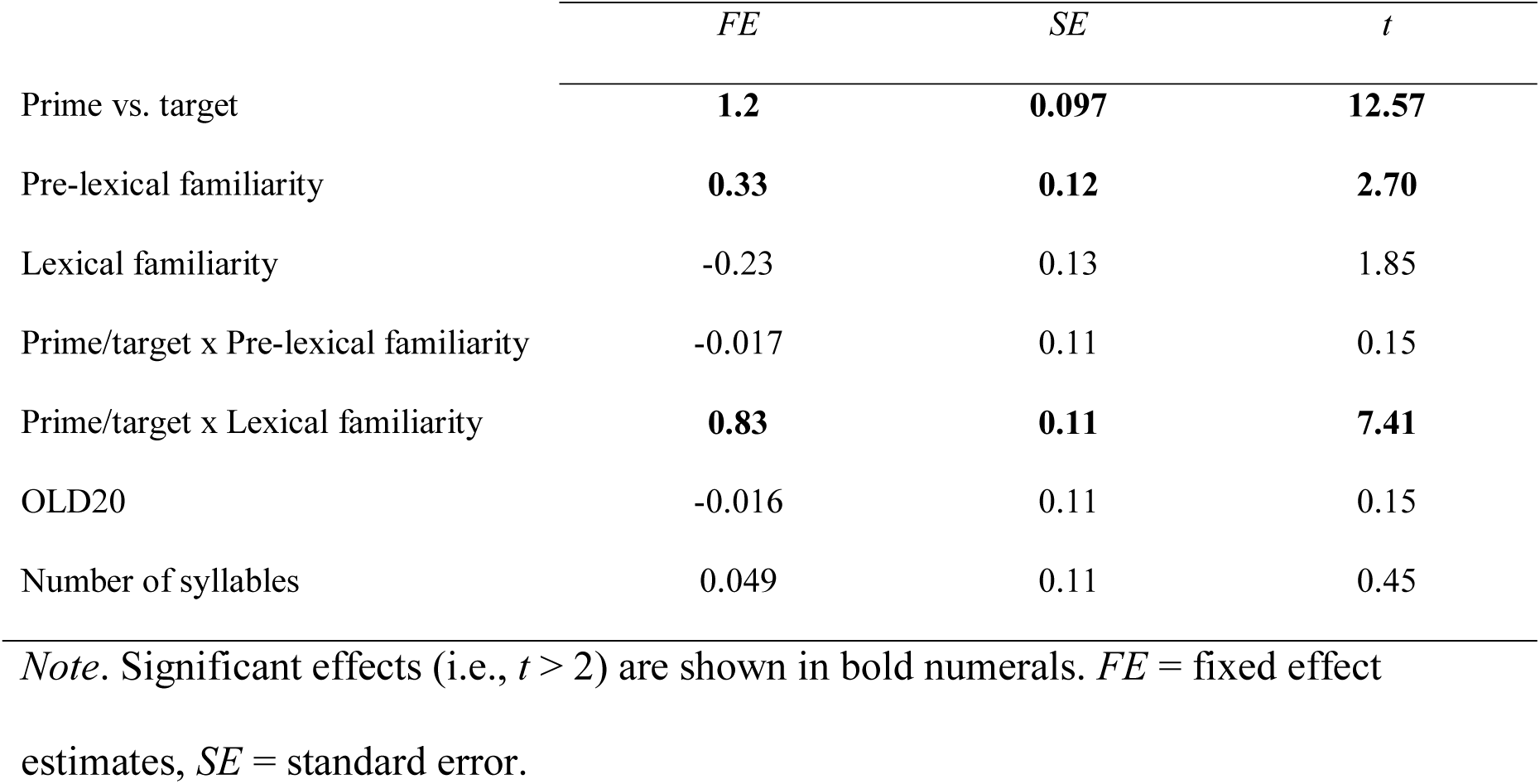
Results from post hoc linear mixed model analyses on ERF values (in 10^-14^ Tesla) from sensor and time point of the strongest effect for the prime/target x lexical familiarity interaction cluster represented in Fig. 5h-j. Separate post hoc analyses for prime and target stimulus are represented in Extended Data Table 5-1.

|  | *FE* | *SE* | *t* |
| --- | --- | --- | --- |
| Prime vs. target | **1.2** | **0.097** | **12.57** |
| Pre-lexical familiarity | **0.33** | **0.12** | **2.70** |
| Lexical familiarity | -0.23 | 0.13 | 1.85 |
| Prime/target x Pre-lexical familiarity | -0.017 | 0.11 | 0.15 |
| Prime/target x Lexical familiarity | **0.83** | **0.11** | **7.41** |
| OLD20 | -0.016 | 0.11 | 0.15 |
| Number of syllables | 0.049 | 0.11 | 0.45 |

*Note.* Significant effects (i.e., $t > 2$ ) are shown in bold numerals. *FE* = fixed effect estimates, *SE* = standard error.

#### Control analyses

To assess potential influences from low-pass filtering, we performed an additional control analyses with a 40 Hz low-pass filter. Results did not differ qualitatively from the results described above obtained with a 20 Hz low-pass filter (Extended Data Fig. 5-4).

We also evaluated the robustness of effects against different choices of baselines by interpreting only those clusters that were significant when prime and target activation were both corrected with the baseline before the prime instead of separate baselines. In addition, we performed a peak-to-peak analysis (cf. Methods section for further details). Significant results from the peak-to-peak analysis strongly support the interaction between prime/target and familiarity effects at left frontal sensors, the main effects of familiarity at left posterior and left frontal sensors, as well as main effects of prime vs. target at bilateral frontal sensors, resembling the effects of clusters CxF, F1, F2, and C3 in Fig. 5 and Extended Data Fig. 5-4 (see Extended Data Fig. 5-1 and 5-5, including also further clusters from the peak-to-peak analysis). Due to the high similarity between standard baseline corrected ERF analysis and peak-to-peak analysis, we conclude that the presented results can be reproduced with a different analysis strategy and therefore are not artificially introduced by the specific choice of baseline correction.

### 2.3 Discussion Experiment 1

The main finding from Experiment 1 (MEG study) was that only lexical familiarity interacted with context (here implemented as contrast prime vs. identical target) to facilitate visual word recognition at the N400 window only. Please note that in the following, for the sake of brevity, we will subsume the processing of words and pseudowords under the term visual word recognition, as we assume similar pre-lexical processing for words and novel pseudowords reflecting the orthographic familiarity (OLD20) match. The finding of stronger repetition suppression for words was consistent with previous studies (Almeida and Poeppel, 2013; Fiebach et al., 2005; but see Deacon et al., 2004; Laszlo and Federmeier, 2007; Laszlo et al., 2012) and identified sources of the effect were compatible with previous localizations of the N400 within the anterior temporal cortex (e.g., Lau et al., 2013a; 2016; Lau and Nguyen, 2015). In contrast, we could not identify a pre-lexical modulation (i.e., an increased reduction of activation for familiarized pseudowords) at any time window. However, we observed a more negative-going amplitude for familiarized PW in contrast to novel PW and words, which is indicative of a pre-lexical familiarity influence irrespective of context. This indicates that our explicit manipulation of pre-lexical familiarity was successful. Also, we found that the pre-lexical familiarity effect was present at the end of the expected time window (e.g., Barber and Kutas, 2007) and localized to the left angular/ supramarginal gyrus, indicating phonological processing; Carreiras et al, 2014; Taylor et al., 2013). We also found strong context effects without modulation of pre-lexical and lexical familiarity. Earliest the effect was present around 100 ms in the occipital cortex. The first presentation elicited a much stronger N100 response compared to the second presentation in repetition trials. In sum, we found an interaction of lexical familiarity and context in the N400 time window but only a main effect of pre-lexical processing.

The interaction of lexical familiarity and context at the N400 indicated within-level context-based facilitation at the lexical level. We related this interaction to lexical level processing since words, i.e., meaningful stimuli, differentiated from pseudowords, i.e., meaningless stimuli. This is in line with previous work (i.e., priming or sentence paradigms: Halgren et al., 2002; Helenius et al., 1998; Lau et al., 2009; 2013a; Rugg, 1985; Simos et al., 1997; Vartiainen et al., 2009) associating this time window and brain locations with lexical-semantic processing. Within-level facilitation was indicated by the finding that the interaction was selective for the N400 time window without indications of interactions at previous time windows. Finally, regarding the mechanistic implementation, the interaction pattern was in line with the expectation of fatigue and predictive coding (see Fig. 1c). This is as sharpening proposes a suppression of the noise in the neuronal signal. Therefore, the difference between words and pseudowords should be easier to detect at the target (i.e., stronger difference; Kok, et al., 2012), which was not the case.

Nevertheless, due to the nature of the task in Experiment 1, we could not investigate whether familiarity-and context-based brain activation differences translate to behavior. Also, we cannot rule out one further potentially confounding influence, i.e., that words and pseudowords differ not only in the availability of lexical-semantic knowledge but also qualitatively with respect to their word status. I.e., even though the familiarity of some pseudowords was temporarily enhanced by the training procedure, the expertise with actual words may be qualitatively different. Moreover, the new experiment included a repetition probability manipulation (i.e., the likelihood of prime and target being the same letter string) that would allow differentiating between fatigue and predictive coding, since only the latter would predict a systematic influence of repetition probability across trials. Finally, in Experiment 1, we were surprised that no evidence for an influence of pre-lexical familiarity on context-based facilitation was found. To replicate the MEG pattern in behavior and systematically examine the role of word status and implement a repetition probability manipulation, we ran a second, behavioral, repetition priming experiment.

## 3 Experiment 2

In Experiment 2, we implemented the explicit investigation of word status by adding a third group of pseudowords. With a paired association task, we associated semantic content to these non-words. Similar as in Experiment 1, we also included familiar pseudowords without meaning. Note, pseudowords with and without semantic associations were visually/perceptually familiarized to the same degree. Therefore, the two groups of pseudowords only varied regarding their associated semantic meaning. Including this additional lexical familiarity condition allowed us to examine potentially different roles of word status and the presence of semantic associations.

We measured behavioral response times in a repetition priming paradigm. Participants had to indicate whether or not a letter string had a semantic association. A yes response would be valid for words and familiarized pseudowords with semantic associations, but not for novel and only perceptually familiarized pseudowords.

As stated above, we also implemented a repetition probability manipulation (i.e., the likelihood of prime and target being the same letter string). Repetition probabilities varied across three blocks, to investigate if the priming effect (i.e., faster responses for repeated vs. non-repeated targets) increases when the local task context allows predicting that the prime is highly likely identical to the target. In previous studies with different visual stimuli higher repetition probability enhanced priming effects, mainly supporting predictive coding (e.g., Grotheer and Kovács, 2014; Olkkonen et al., 2017; Summerfield et al., 2008; 2011).

### 3.1 Methods

#### 3.1.1 Participants

24 healthy native speakers of German recruited from university campuses (16 females, mean age 23.1±3.4 years, range 19-31 years, 22 right-handers) were included in the final data analysis. All participants had normal or corrected-to-normal vision, and normal reading abilities as assessed with the adult version of the Salzburg Reading Screening (unpublished adult version of Mayringer and Wimmer, 2003). Further participants were excluded at different stages of the experiment due to the following reasons: Low reading skills (i.e., reading test score below 16^th^ percentile; N = 4), insufficient performance during pseudoword familiarization (i.e., accuracy for semantic or familiar pseudowords < 50 % in the final learning session; N = 3), or failure to complete the experimental protocol (N = 2). Four participants were excluded after data analysis due to insufficient performance (< 25 % correct for non-repeated words). All participants gave written informed consent according to procedures approved by the local ethics committee and received 10 € per hour or course credit as compensation.

#### 3.1.2 Stimuli and presentation procedure

60 German nouns (half natural and half man-made; logarithmic word frequency: mean ± standard error = 1.93 ± 0.09, range 0.00 to 3.30) and 180 pronounceable pseudowords with characteristics similar to Experiment 1 were presented in a repetition priming task. Pseudowords were divided into three sets, each of which was matched to the word set on orthographic similarity (OLD20, Yarkoni et al., 2008; words: 1.538 ± 0.038; pseudowords: 1.605 ± 0.032, 1.542 ± 0.045, and 1.596 ± 0.044) and number of syllables (1.833 ± 0.059; 1.95 ± 0.028; 1. 967 ± 0.023; 1.9 ± 0.039, respectively; Table 1). Participants were perceptually familiarized with one set analogous to Experiment 1, and additionally learned semantic associations for a second set within a paired-association task (see below for details). The third set of pseudowords was never seen by the participants before the priming task. For the familiarization procedure, two sets of 60 object images each were chosen from the Bank of Standardized Stimuli (BOSS; Brodeur et al., 2010; 2014) such that German object names assigned to the images were matched between the two sets for logarithmic word frequency (set means: 2.093 ± 0.081; 2.070 ± 0.077), OLD20 (set means: 1.639 ± 0.054; 1.630 ± 0.053), and number of syllables (set means: 2.000 ± 0.071; 2.000 ± 0.071). Object names were determined by having four independent participants write down for each object the name they considered most suitable; only objects for which at least three participants provided the same name were selected. The two sets of object images finally selected were matched on available ratings of familiarity (set means: 4.364 ± 0.040; 4.333 ± 0.043), object agreement (i.e., rated similarity between an object imagined by the participants upon perceiving the object’s name, and the actual object image; set means: 3.910 ± 0.056; 3.901 ± 0.064), and rated subjective visual complexity (set means: 2.426 ± 0.058; 2.475 ± 0.066; Brodeur et al., 2014), analogous to procedures reported by Breitenstein et al. (2007).

Six variants of the familiarization task were prepared, across which the assignment of the three pseudoword sets to the familiarized, i.e., familiar vs. semantic, as well as to the novel condition was varied (see Table 6). In addition, the assignment of the two object image sets to the familiarized pseudowords with and without semantic associations was varied. Note that for 18 of the 24 participants, the six experimental versions, as well as the order of blocks and response hands in the repetition priming task (see below), were counterbalanced. In addition, six participants were included from the pilot investigation in which this was not the case (all had the same response hands and the initial block had a repetition probability of 25 %). Results did not differ qualitatively when these participants were included or not. Stimulus presentation procedures were identical to those of behavioral sessions of Experiment 1 (Fig. 2a), with the exception that the background was set to grey.

**Table 6.** Overview of the six experimental versions A to F of Experiment 2, indicating which of the two object image sets (A, B) was learned and which of the three pseudoword sets (1, 2, 3) was assigned to which familiarity condition.

| Version | Learned object set | Pseudoword set |  |  | Participants |
| --- | --- | --- | --- | --- | --- |
|  |  | *Semantic* | *Familiar* | *Novel* |  |
| A | A | 1 | 2 | 3 | 8 |
| B | B | 2 | 1 | 3 | 4 |
| C | A | 2 | 3 | 1 | 3 |
| D | B | 3 | 2 | 1 | 3 |
| E | A | 3 | 1 | 2 | 3 |
| F | B | 1 | 3 | 2 | 3 |

*Note.* Novel = pseudowords first shown in the repetition priming task. Participants refers to the number of participants assigned to each version.

#### 3.1.3 Pseudoword Familiarization

Participants performed five pseudoword familiarization sessions in the course of three consecutive days, i.e., two sessions each on day 1 and 2, and one session on day 3 (before the repetition priming task). Each session lasted about one hour, and participants could take a short break after the first half, as well as a mandatory one-hour break before the next session. Each session consisted of reading aloud each pseudoword (mean error rate across sessions: 1.4 %), a computer-based paired-association task with congruent vs. incongruent parings of pseudowords and object images, and a naming task. While one set of pseudowords was familiarized pre-lexically as in Experiment 1, i.e., merely through repeated exposure (‘familiar pseudowords’), one set was additionally associated with semantic information (‘semantic pseudowords’). The paired-association procedure was adapted from previous studies (e.g., Breitenstein and Knecht, 2002; Breitenstein et al., 2007; Dobel et al., 2009), however using visual instead of auditory pseudowords and naturalistic photographs of objects instead of line drawings (see above). Furthermore, we used an explicit instead of an implicit learning instruction in order to establish strong associations between pseudowords and the assigned meanings.

Familiar and semantic pseudowords were presented in random order for 800 ms, followed by an object image (horizontal and vertical visual angles 15.8°) for 1,500 ms or until response (Fig. 2c). During the ITI of 1,000 ms, two vertical black bars indicating the center of the screen where participants were asked to fixate were presented. Each pseudoword was presented four times in the first and four times in the second half of each session (960 trials in total per session). Semantic pseudowords were arbitrarily but above-chance (i.e., six out of eight presentations) matched with object images so that participants could learn to associate their meaning over the course of the familiarization sessions. This ratio was chosen so that despite successful learning, false alarms could be investigated which provide important information on participants’ sensitivity. In contrast, familiar pseudowords were followed by a different object image in each trial.

Participants were asked to learn a meaning for the presented pseudowords based on the frequency with which the pseudowords were paired with certain object images. They were explicitly informed about the inconsistent pairings for half of the pseudowords. Participants were instructed to silently read the presented pseudowords and to respond as accurately and quickly as possible, whether a presented object image matched the preceding pseudoword or not. In addition, they were encouraged to guess if insecure. Participants responded by pressing one of two buttons on a keyboard with either the left or right index finger. To prevent potential response biases, the assignment of response hand and response varied from trial to trial (by presenting a red bar indicating non-match on one side and a green bar indicating match on the other side of the object image). In the first familiarization session, participants completed a short practice block of ten trials before the start of the actual paired-association task.

In the naming task (Fig. 2d), each pseudoword from the paired-association task was presented once. Participants were instructed to name its associated object if an association could be retrieved, or to respond “weiter” (German for “next”) whenever this was not possible. The experimenter wrote down the participants’ responses and logged the three possible responses (correct, incorrect, next) into the presentation software. Responses were considered correct whenever a name suitable for the corresponding object was provided (e.g., “cabin” instead of “barn”). Participants did not receive feedback.

LMMs (including participant, object image and pseudoword as random effects on the intercept; see Experiment 1 Methods) revealed that d’ for the paired-association task significantly increased across sessions from 0.45 in session 1 to 2.04 in session 5 (main effect of session: *estimate* = 0.54, *SE* = 0.035, *t* = 15.70; Fig. 3c; see Extended Data Fig. 3-1 including post hoc analyses for pairwise sessions). This indicates that participants improved in identifying matching and non-matching pseudoword-object combinations. In the final familiarization session, participants reached high mean accuracies of 92.68 % (range: 74.72 to 99.72) for the identification of matching objects for semantic pseudowords, and 89.84 % (range: 60.47 to 99.38) for the identification of non-matching objects for pseudowords familiarized without semantics (Fig. 3d). Importantly, participants also demonstrated high average accuracies of 95.68 % (range: 88.14 to 100) for semantic pseudowords in case they were presented with a non-matching object (Fig. 3d), indicating that their high performance for matching pseudoword-object combinations cannot be attributed to a response bias, i.e., responding “match” whenever a semantic pseudoword was presented.

In the pseudoword naming task, which was administered at the end of each familiarization session, LMMs (including participant and item as random effects on the intercept) revealed that d’ significantly increased from 0.23 in session 1 to 2.43 in session 5 (main effect of session: *estimate* = 0.77, *SE* = 0.040, *t* = 19.41; Fig. 3e; see Extended Data Fig. 3-1 including post hoc analyses for pairwise sessions). In the final session, participants named the correct object for between 51.67 to 100 % of semantic pseudowords (mean 78.61; Fig. 3f, left) and refrained from a response for 61.67 to 100 % of pseudowords familiarized without semantics (mean 90.28; Fig. 3f, right), indicating that they indeed learned the corresponding meaning for semantic pseudowords.

#### 3.1.4 Repetition Priming

Following the fifth familiarization session on day 3, participants completed a repetition priming experiment after a break of at least one hour. Experimental procedures were analogous to those described for Experiment 1, with the following exceptions: Semantic pseudowords were presented as additional familiarity condition, and no catch trials were presented. The prime stimulus in each trial was preceded by 800 ms of hash mark presentation. The inter-trial interval varied between 800 and 1,200 ms. Furthermore, the repetition probability was varied across the three experimental blocks. 15 participants first completed a block with 25 % repetition probability, followed by 50 % in the second and 75 % in the last block; the remaining nine participants completed the blocks in the reverse order. Participants were informed about the repetition probabilities at the start of each block. Their task was to silently read the presented letter strings and respond as accurately and quickly as possible to the second letter string in each trial, whether they could explicitly associate a meaning or not (button presses on a keyboard with left/right index finger; dominant vs. non-dominant hand for yes-response: 13 vs. 11 participants, respectively). This task was chosen to elicit the same response for semantic pseudowords as for words. Each letter string (i.e., word or pseudoword) was presented once per block, either in the repetition or in the non-repetition condition. In total, 240 trials (60 per condition) were presented in each block. Letter strings were used at maximum twice for non-repetition trials; in this case, they were combined with two different letter strings. Prior to the task, eight practice trials were completed. The total duration of the priming task was around 45 min.

#### 3.1.5 Analyses

Analogous to the analysis of the pseudoword familiarization procedure, behavioral data of the repetition priming task were analyzed using LMMs allowing random effects of both participant and items (prime and target stimulus) on the intercept, as well as analysis of imbalanced data (Baayen et al., 2008). We mainly focused on response times of correct responses, but also investigated accuracies using generalized LMMs with a binomial link function. Response times were log transformed to account for their skewed ex-Gaussian distribution.

We first performed an analysis with factors repetition congruency (repetition vs. non-repetition trials), reflecting the main manipulation of context effects in Experiment 2, and repetition probability (25, 50 or 75 %). To assess pre-lexical and semantic contributions to behavioral context and familiarity effects, we investigated the four-way interaction between repetition congruency, repetition probability, pre-lexical, and lexical familiarity. The latter two were manipulated orthogonally, such that familiarity was entered as two factors coding pre-lexical (0: novel pseudowords and words; 1: familiar pseudowords with and without semantics) and lexical familiarity (0: novel and familiar pseudowords without semantics; 1: semantic pseudowords and words). Since context provided by the prime stimulus might override familiarity effects (Kretzschmar et al., 2015), we additionally investigated the three-way interaction between pre-lexical familiarity, lexical familiarity, and repetition probability in non-repetition trials only (i.e., in the absence of valid contextual information). Note that for repetition priming analyses, we set behavioral responses from the first block (i.e., with 75 % repetition probability) of one participant to NA, because she reported a misinterpretation of the task instruction that was clarified for the final two blocks.

All (generalized) LMMs included the interactions of all fixed effects described so far. Since not all trials entered the analyses (due to miss trials and for the response time analysis due to exclusion of trials with incorrect responses), which might have affected the match across letter string conditions, OLD20 and number of syllables were included as additional fixed effects. All fixed effects were centered and z-transformed. For each significant interaction, pairwise differences between conditions were investigated using post hoc linear mixed models including only the relevant conditions.

### 3.2 Results

In the semantic association judgments of the repetition priming experiment, average accuracies for repetition trials were high, albeit not at ceiling, across all repetition probabilities (86.9 %, 85.8 %, and 83.9 % for 75 %, 50 %, and 25 % repetition probability, respectively; Extended Data Fig. 6-1a), as well as across all familiarity conditions with the exception of familiarized pseudowords with semantic associations (90.7 %, 88.1 %, 72.2 %, and 91.1 % for novel pseudowords, familiarized pseudowords without and with semantic associations, and words, respectively; Extended Data Fig. 6-1b). The lower accuracy for semantic pseudowords indicates that participants did not establish a semantic association with all (but yet the majority of) pseudowords, which is also consistent with their performance in the final naming session (see Analyses section and Fig. 3f). As a consequence, we only used correct trials for the response time analysis. Accuracies in non-repetition trials were overall lower (82.6 %) compared to repetition trials (88.5; Extended Data Fig. 6-1a). Statistical analyses of accuracies can be found in Extended Data Tables 10-1 and 10-2. In contrast to the MEG analysis, we included familiarity effects related to pre-lexical and lexical familiarity as two separate factors, since pre-lexical familiarity was manipulated orthogonally to lexical familiarity (cf. Methods). In the following, we report the effects on response times most relevant for our hypotheses, while Tables 7 to 10 provide a detailed overview of all statistical results.

**Table 7.**
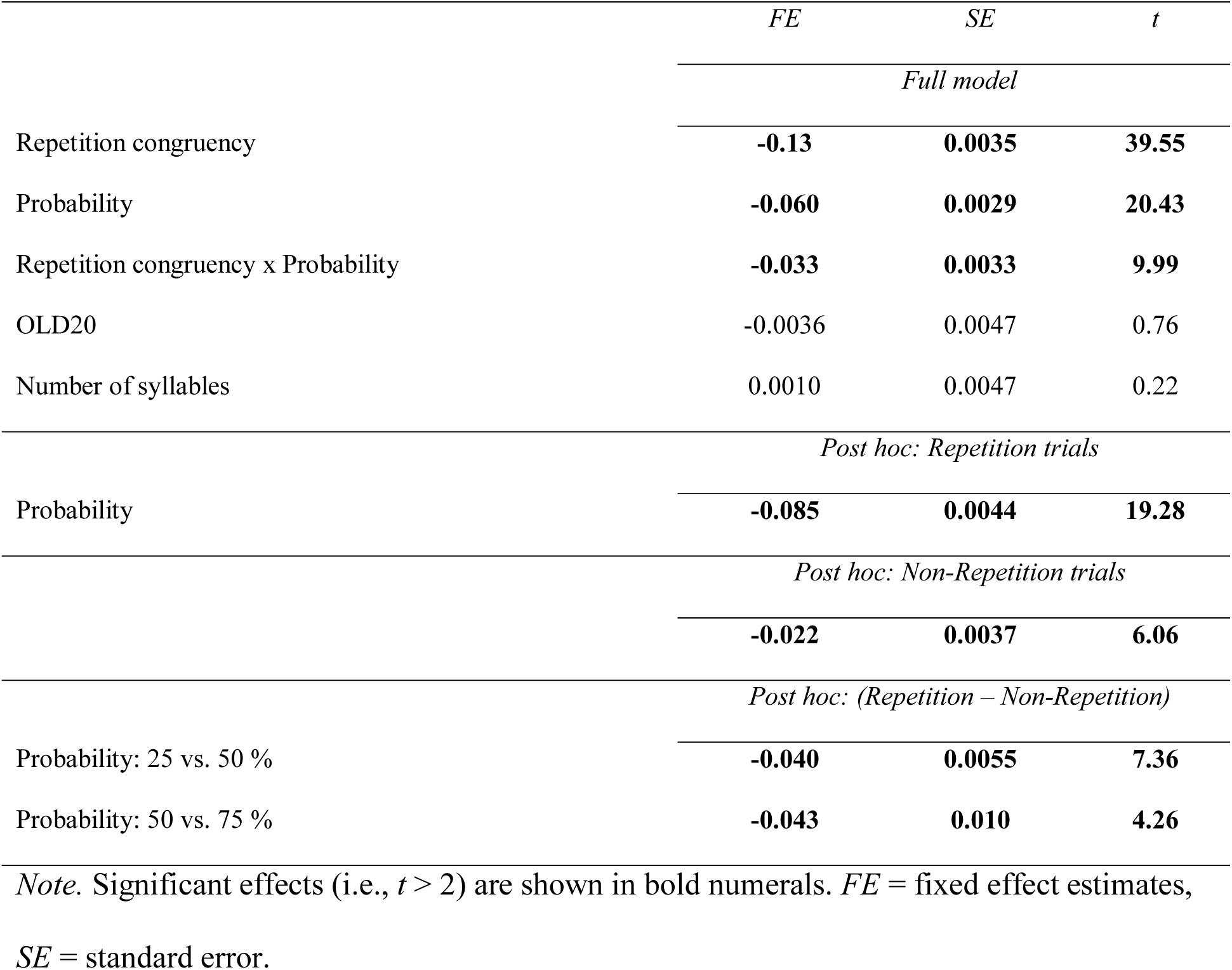
Results from the linear mixed model analyses investigating repetition congruency (repetition vs. non-repetition) and repetition probability in log transformed response times from the repetition priming task (Experiment 2).

#### Repetition priming

For a first manipulation check we investigated the influence of repetition probability on context effects irrespective of familiarity conditions (Fig. 6a and statistics in Table 7). Here, context effects refer to the classical priming effect reflected in the repetition congruency contrast (repetition vs. non-repetition). Response times showed a significant interaction between this priming effect and repetition probability. The interaction revealed a decrease in response times with increasing repetition probability (main effect of repetition probability: *estimate* = −0.060, *SE* = 0.0029, *t* = 20.43) which was stronger for repetition (*estimate* = −0.085, *SE* = 0.0044, *t* = 19.28) compared to non-repetition trials (*estimate* = −0.022, *SE* = 0.0037, *t* = 6.06). I.e., the priming effect (difference between repetition and non-repetition trials) was smaller for a repetition probability of 25 vs. 50 % (*estimate* = −0.040, *SE* = 0.0055, *t* = 7.36) and smaller for 50 vs. 75 % (*estimate* = −0.043, *SE* = 0.010, *t* = 4.26; Fig. 6a; Table 7). This finding indicates that context effects increase when they can be expected more reliably.

**Figure 6.**
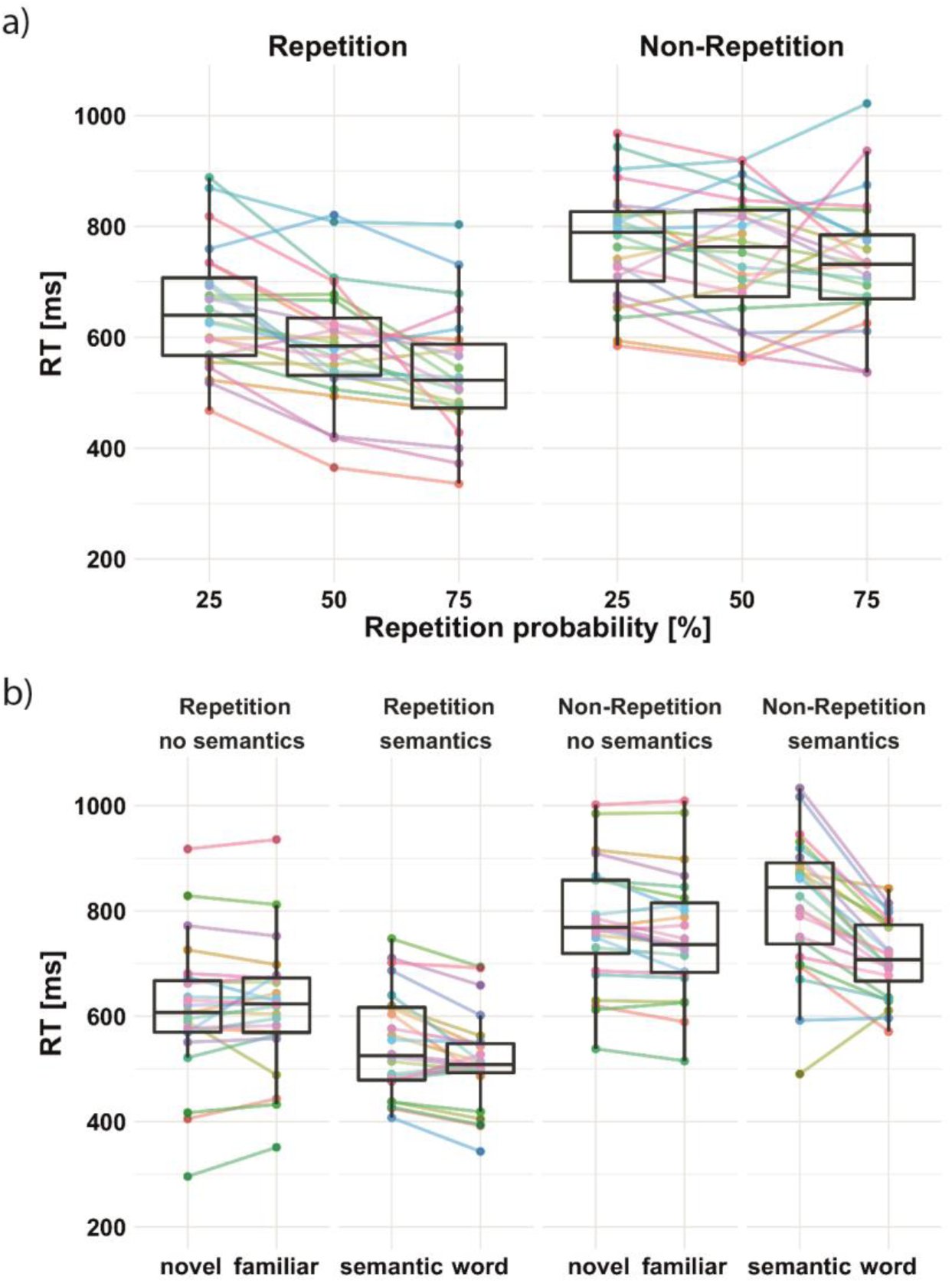
Response times in Experiment 2 (semantic association judgment task). a) Repetition probability effect for repetition (left) and non-repetition trials (right) averaged across familiarity conditions. b) Familiarity influence on context effects, i.e., repeated (left) vs. non-repeated targets (right). Effects are separated for familiarity conditions, with an additional separation for letter strings with and without semantic associations, averaged across repetition probabilities. Colored dots and lines represent individual participants. Accuracy data are shown in Extended Data Fig. 6-1.

#### Familiarity effects

To investigate the influence of pre-lexical and lexical familiarity in the absence of valid contextual information, we focused on non-repetition trials. We observed a significant interaction between pre-lexical and lexical familiarity (Table 8). This interaction was driven by the strong difference in response times for the two semantic letter string groups: Semantic pseudowords showed the longest response times (all t’s > 4 for post hoc contrasts of semantic pseudowords vs. the other three conditions; see Table 9 for details), reflecting the specific difficulty of retrieving semantics for a newly acquired vocabulary, particularly in case of unfulfilled expectations. This notion is also in line with the accuracy data (see Extended Data Fig. 6-1b). However, faster response times for words compared to novel (*estimate* = −0.039, *SE* = 0.0064, *t* = 6.06) and familiar pseudowords (*estimate* = −0.022, *SE* = 0.0063, *t* = 3.52; Table 9) indicate facilitated processing of letter strings with both fully established semantic associations and word status. In addition, response times were faster for familiar vs. novel pseudowords (*estimate* = −0.016, *SE* = 0.0046, *t* = 3.49).

**Table 8.**
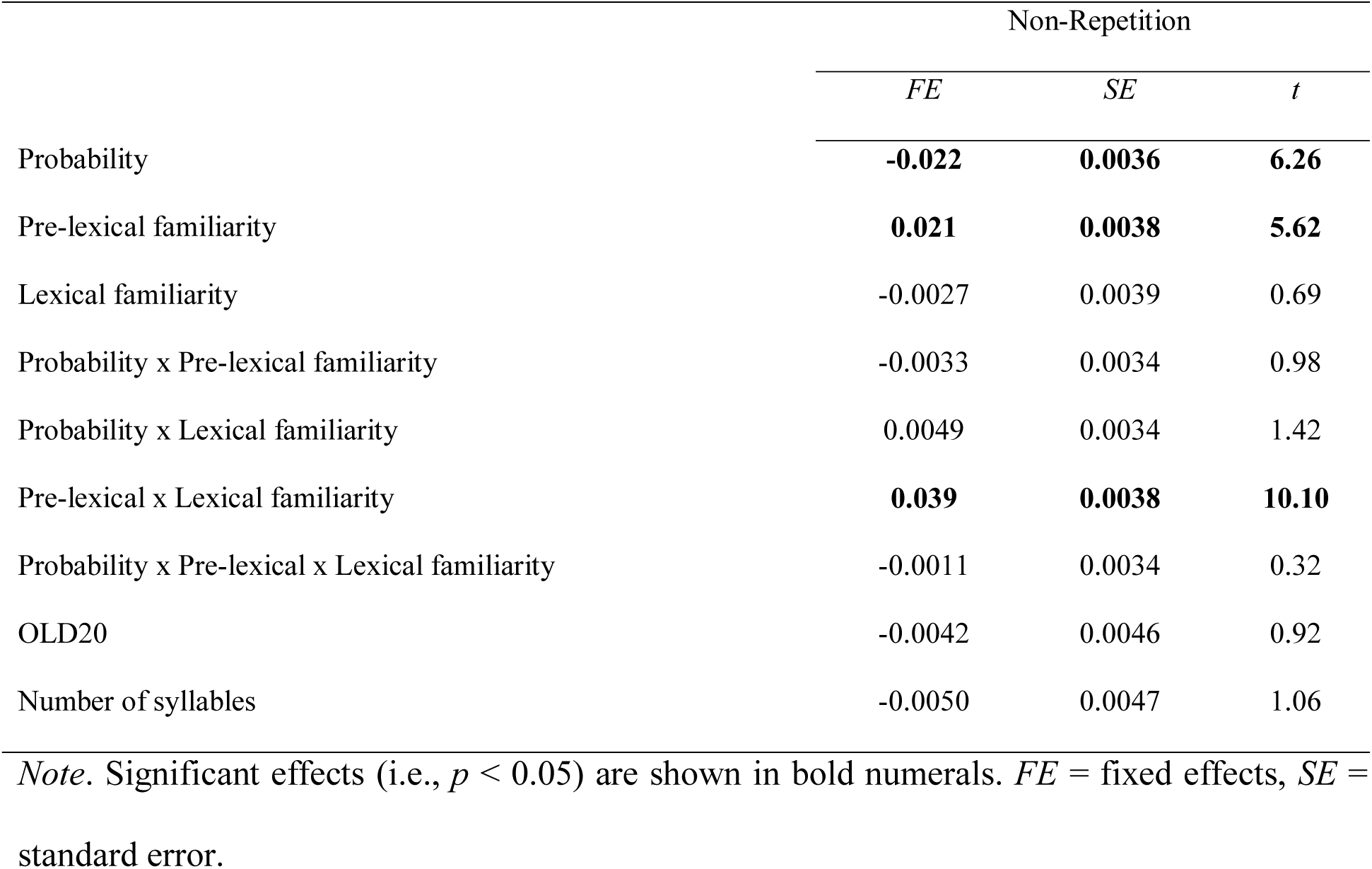
Results from the linear mixed model analyses investigating repetition probability, familiarity, and semantics in log transformed response times from non-repetition trials only (Experiment 2).

**Table 9.** Results from post hoc linear mixed model analyses investigating familiarity effects in log transformed response times in the repetition priming task (Experiment 2).

|  | Repetition |  |  | Non-Repetition |  |  |
| --- | --- | --- | --- | --- | --- | --- |
|  | *FE* | *SE* | *t* | *FE* | *SE* | *t* |
| Novel vs. familiar PW |  |  |  |  |  |  |
| Familiarity | 0.0058 | 0.0056 | 1.03 | **-0.016** | **0.0046** | **3.49** |
| OLD20 | -0.016 | 0.0085 | 1.91 | -0.0053 | 0.0058 | 0.91 |
| Number of syllables | -0.0077 | 0.0082 | 0.94 | -0.0083 | 0.0059 | 1.39 |
| Novel vs. semantic PW |  |  |  |  |  |  |
| Familiarity | **-0.048** | **0.0059** | **8.11** | **0.023** | **0.0051** | **4.54** |
| OLD20 | -0.012 | 0.0075 | 1.56 | -0.011 | 0.0064 | 1.68 |
| Number of syllables | -0.0085 | 0.0074 | 1.14 | **-0.015** | **0.0067** | **2.16** |
| Novel PW vs. words |  |  |  |  |  |  |
| Familiarity | **-0.076** | **0.0068** | **11.24** | **-0.039** | **0.0064** | **6.06** |
| OLD20 | -0.016 | 0.0067 | 2.39 | -0.0042 | 0.0061 | 0.69 |
| Number of syllables | 0.0031 | 0.0069 | 0.45 | 0.0022 | 0.0067 | 0.32 |
| Familiar vs. semantic PW |  |  |  |  |  |  |
| Familiarity | **-0.051** | **0.0060** | **8.52** | **0.039** | **0.0053** | **7.25** |
| OLD20 | -0.0056 | 0.0077 | 0.72 | -0.0064 | 0.0063 | 1.03 |
| Number of syllables | -0.0066 | 0.0076 | 0.86 | **-0.014** | **0.0064** | **2.22** |
| Familiar PW vs. words |  |  |  |  |  |  |
| Familiarity | **0.080** | **0.0079** | **10.08** | **-0.022** | **0.0063** | **3.52** |
| OLD20 | -0.0087 | 0.0077 | 1.12 | 0.0007 | 0.0060 | 0.12 |
| Number of syllables | 0.0049 | 0.0081 | 0.60 | -0.0008 | 0.0066 | 0.12 |
| Semantic PW vs. words |  |  |  |  |  |  |
| Familiarity | **0.026** | **0.0071** | **3.64** | **0.064** | **0.0073** | **8.74** |
| OLD20 | -0.0030 | 0.0070 | 0.43 | -0.0078 | 0.0070 | 1.11 |
| Number of syllables | 0.0038 | 0.0074 | 0.52 | -0.0055 | 0.0078 | 0.70 |

Note. Significant effects (i.e., $t > 2$ ) are shown in bold numerals. *FE* = fixed effects,
*SE* = standard error.

#### Influence of pre-lexical and lexical familiarity on context effects

Repetition probability did not interact with pre-lexical or lexical familiarity (all *t*’s < 1, including the 3-way interaction; see Table 10). However, a significant interaction between repetition congruency and lexical familiarity revealed stronger priming effects for letter strings with semantic associations (i.e., words and semantic pseudowords) in comparison to pseudowords without semantic associations (*estimate* = −0.033, *SE* = 0.0030, *t* = 11.13; Table 10). In repetition trials, the response times for pseudowords with associated semantics were lower than for the other pseudoword conditions (pairwise post hoc contrasts: semantic vs. novel pseudowords: *estimate* = −0.048, *SE* = 0.0059, *t* = 8.11; semantic vs. familiar pseudowords: *estimate* = −0.051, *SE* = 0.0060, *t* = 8.52; Table 9). This indicates that the involvement of semantic information increases context effects dramatically, even reversing familiarity effects found in the absence of context-based facilitation.

**Table 10.** Results from the linear mixed model analyses investigating repetition congruency (repetition vs. non-repetition), probability, familiarity, and semantics in log transformed response times in the repetition priming task (Experiment 2). Statistical analyses of accuracy data can be found in Extended Data Tables 10-1 and 10-2.

|  | *FE* | *SE* | *t* |
| --- | --- | --- | --- |
| Repetition congruency | **-0.13** | **0.0031** | **41.76** |
| Probability | **-0.060** | **0.0029** | **20.53** |
| Pre-lexical familiarity | **0.019** | **0.0032** | **5.90** |
| Lexical familiarity | **-0.036** | **0.0033** | **10.80** |
| Repetition congruency x Probability | **-0.032** | **0.0031** | **10.29** |
| Repetition congruency x Pre-lexical familiarity | -0.0026 | 0.0029 | 0.88 |
| Repetition congruency x Lexical familiarity | **-0.033** | **0.0030** | **11.13** |
| Probability x Pre-lexical familiarity | -0.0036 | 0.0029 | 1.26 |
| Probability x Lexical familiarity | 0.0057 | 0.0029 | 1.98 |
| Pre-lexical x Lexical familiarity | **0.026** | **0.0032** | **7.86** |
| Repetition congruency x Probability x Pre-lexical familiarity | 0.00045 | 0.0029 | 0.15 |
| Repetition congruency x Probability x Lexical familiarity | 0.00065 | 0.0029 | 0.22 |
| Repetition congruency x Pre-lexical x Lexical familiarity | **-0.013** | **0.0030** | **4.47** |
| Probability x Pre-lexical x Lexical familiarity | -0.0020 | 0.0029 | 0.69 |
| Repetition congruency x Probability x Pre-lexical x Lexical familiarity | -0.00054 | 0.0029 | 0.18 |
| OLD20 | -0.0065 | 0.0036 | 1.80 |
| Number of syllables | -0.0052 | 0.0037 | 1.42 |

*Note.* Significant effects (i.e., $p < 0.05$ ) are shown in bold numerals. *FE* = fixed effects, *SE* = standard error.

### 3.3 Discussion Experiment 2

In general, the behavioral results replicated the main MEG findings. Context-based facilitation, here implemented as repetition congruency, on response times was stronger for words and pseudowords with semantic associations in contrast to pseudowords without meaning. The interaction of lexical familiarity and context replicated the modulation of lexical information on context-based facilitation found in the N400. We also observed that recently increased pre-lexical familiarity, in familiarized pseudowords without meaning, resulted in faster response times compared to novel pseudowords in non-repetition trials, i.e., in the absence of valid contextual information. This finding is compatible with the increased activation to familiar pseudowords within the left angular/ supramarginal gyrus shown in the MEG. Finally, we found strong general priming effects that replicate the strong context effects demonstrated in the MEG results. Once more, we found no evidence for an interaction of pre-lexical familiarity and context-based facilitation.

A word status effect, i.e., words versus pseudowords with associated meaning, was also found in response times, but the difference in response times was much smaller when the contextual information was valid. This pattern indicates that even recently learned letter strings use semantic meaning to facilitate word recognition on a lexical level in a predictable context (cf. Tamminen and Gaskell, 2013; van der Ven et al., 2015). Also, we could identify a strong repetition probability effect. Here a higher repetition probability resulted in faster response times in repetition trials replicating previous studies with different visual stimuli (Barbosa and Kouider, 2018; Olkkonen et al., 2017). We expected this finding when one implements a predictive coding mechanism. Thus, fatigue is ruled out as a mechanism for context-based facilitation.

## 4 Discussion

In the two experiments of the present study, we found evidence for context-based facilitation of visual word recognition within the lexical-semantic processing level based on a predictive coding mechanism. Most prevalent was the increased facilitation, reflected in reduced brain activation at the left anterior temporal cortex around 400 ms and faster behavioral responses, when semantic information was present (see Fig. 7). We found no evidence for context-based facilitation through pre-lexical (i.e., orthographic and/or phonological) familiarity. Also, we could not detect evidence for top-down facilitation, as there was no influence of lexical information on context effects in earlier time windows (i.e., < 400 ms) associated with visual and pre-lexical processing. At the level of visual processing, we found familiarity-unspecific repetition effects in the occipital cortex around 100 ms. Combined, we take this pattern as evidence for context-based facilitation within lexical and visual processing levels (Fig. 7).

**Figure 7.**
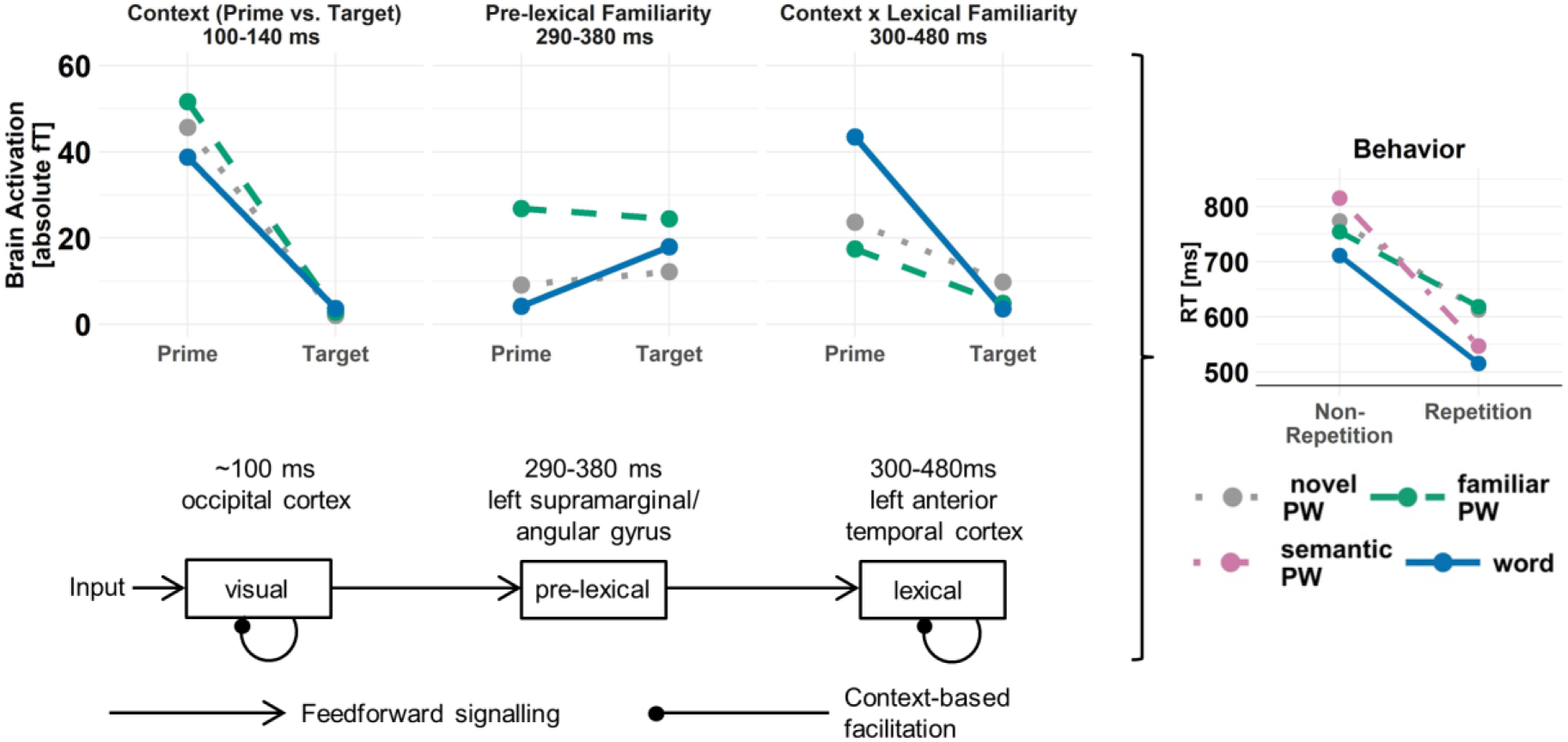
Summary of our findings (top) and implications for the comprehensive connectionist model of visual word recognition (cf. Carreiras et al., 2014; bottom). In the upper part of the figure, we present MEG findings for each processing level separated for the letter string conditions (words and pseudowords: PW). On the right, we present the behavioral pattern from response times. Note, all data figures are presented in a simplified form (combining individual data points and excluding participant outliers outside of 1.5 times the interquartile range above the upper quartile and below the lower quartile) allowing a more lucid presentation of identified effect patterns. For more details see Fig. 5 for the MEG data and Fig. 6 for behavior. Our evidence supports context-based facilitation within visual and lexical processing levels (bottom).

### Implications for neuro-cognitive models of visual word recognition

One of the main goals of this study was to investigate whether context-based facilitation of visual word recognition is implemented via top-down feedback from higher to lower processing levels, or restricted to within each processing level: i.e., assessing the architecture of context-based facilitation. The two alternative architectures were formalized in previous neuro-cognitive models of visual word recognition. Laszlo and Amstrong (2014) implemented a strictly feed-forward model including within-level context-based facilitation. The model architecture brought forward by Carreiras et al. (2014) additionally included top-down connections. The present evidence favors the architecture implemented by Laszlo and Amstrong (2014). As described above, at the lexical processing level we could find within-level context-based facilitation but no evidence for cross-level top-down facilitation. The latter is a central component of the architecture described by Carreiras et al., (2014).

Despite not finding evidence for pre-lexical influences on context effects in the present study, we could identify an activation cluster in the left angular/ supramarginal cortex, showing a pre-lexical familiarity effect. At the sensor level, the activation in response to familiarized pseudowords differed from novel pseudowords and words without an interaction with context. In behavior, we also observed facilitated recognition of pre-lexically familiarized in contrast to novel pseudowords when no valid contextual information was available. These findings reflect that pre-lexical familiarity has a central role in visual word recognition (see also Bermúdez-Margaretto et al., 2015; Glezer et al., 2015; Xue and Poldrack, 2007), but no evidence for an increased context-based facilitation within the prelexical processing levels, as proposed by Laszlo and Amstrong (2014), was found.

One could argue that the learning paradigm did not build up a sufficiently strong pre-lexical representation to influence context-based facilitation. In our opinion, this is less likely for two reasons. In both experiments, we could find pre-lexical facilitation when no valid context-based expectations could be formed. This finding is a manipulation check showing that pseudoword learning was successful. The second reason is that in the behavioral experiment, the learned lexical-semantic information was successfully utilized to increase context-based facilitation (see Tamminen and Gaskell, 2013; van der Ven et al., 2015 for similar results). Both these findings strongly indicate that the learning paradigms used here were effective in influencing processing of the learned pseudowords. Thus, these findings underline the surprising result that additional pre-lexical familiarity did not increase context-based facilitation. In addition, a previous study found no reliable influence of OLD20 on context effects in sentences (Payne et al., 2015). Thus, learning-independent manipulations of pre-lexical familiarity did also not modulate context-based facilitation.

Previous studies using text-or sentence-based context manipulations found top-down influences on visual or pre-lexical processing (e.g., Brothers et al., 2015; Dambacher et al., 2006; Kim and Lai, 2012; Lee et al., 2012). However, time point (i.e., N170 vs. P2 vs. N200/250 component) and direction of these top-down effects were highly inconsistent (see Nieuwland, 2019 for a review). One limitation of sentence studies might be that word predictability out of a sentence context, reflecting lexical-semantic top-down information, and pre-lexical familiarity (e.g., OLD20) are naturally confounded. For example, in the Potsdam Sentence Corpus used by Dambacher et al. (2006), sentence-level predictability and OLD20 (i.e., item-level word familiarity) correlated with an r of -.24. Typically, these studies control for word characteristics like word frequency (r with predictability:.33), which is also associated with semantic word characteristics (e.g., r with the semantic neighborhood around .75; Goh et al., 2016; Yap et al., 2012), but not orthographic characteristics like OLD20. This confound pattern might indicate that only the combined availability of predictable sentence-level context and high orthographic familiarity enables early context effects, which should be explicitly investigated in future studies.

### Mechanistic implementation of context-based facilitations

As pointed out above, the interaction pattern at the N400 component, in particular, the reduction of the difference of words against pseudowords, was informative to determine that predictive coding was the most probable mechanism underlying the context-based facilitation phenomena. At the prime, words showed a stronger N400 in contrast to pseudowords. At the target, this difference was reversed. This pattern rules out sharpening, as in sharpening the expectation is a suppression of the noise in the neuronal signal (cf. Blank and Davis, 2016; Kok et al., 2012; Richter et al., 2018). Therefore, the difference between words and pseudowords should be easier to detect (i.e., stronger difference). The N400 activation shows the opposite pattern when comparing prime (words > pseudowords) and target amplitude (words < pseudowords; Fig. 7). We consider the change in effect direction as evidence against a sharpening mechanism. Evidence against a fatigue mechanism was the finding that a high repetition probability, across trials, resulted in stronger context effects in response times. This finding was only expected by predictive coding and previous evidence from neuronal (e.g., Delaney-Busch et al., 2017; Grotheer and Kovács, 2014; Lau et al., 2013b; Mayrhauser et al., 2014; Summerfield, 2008; 2011; Todorovic et al., 2011) and behavioral investigations (Barbosa and Kouider, 2018; Olkkonen et al., 2017) came to similar conclusions.

In addition, the interaction between with vs. without context (i.e., prime vs. target) and valid vs. invalid context (i.e., repetition vs. non-repetition) at the N400 also provides evidence against fatigue. A fatigue mechanism cannot explain the increased activation for unexpected targets in non-repetition trials compared to primes. At the same time, this increase fits well with predictive coding. In Experiment 1, the repetition probability was 75%. As a consequence, a repetition was likely expected in every trial. Irrespective of repetition or non-repetition trials, this expectation is transformed in a prediction and, in case of a non-repetition trial, the prediction is not met resulting in a prediction error. The increase in the N400 amplitude for non-repeated targets vs. primes might indicate a higher prediction error for mispredicted vs. unpredicted stimuli (e.g., Hsu et al., 2015; 2018). Thus, these findings indicate that a predictive coding mechanism offers the most appropriate explanation for context-based facilitation described here.

The current interpretations concerning the architecture and mechanistic implementation of context-based facilitation are not necessarily compatible. First, our favored architecture (Laszlo and Armstrong, 2014) implemented a fatigue mechanism. As pointed out above, the expected patterns from fatigue and predictive coding are relatively similar. Only our repetition probability and repetition congruency manipulations allowed the differentiation of predictive coding and fatigue. We expect that the implementation of a fatigue mechanism will not be able to simulate the effect of repetition probability presented here.

Still, another incompatibility is prevalent. We could not find evidence for the assumptions concerning the processing levels involved in context-based facilitation and the architecture proposed by the predictive coding theory (e.g., Friston, 2005; Rao and Ballard, 1999). Predictive coding assumes that, at all processing levels, one integrates all available information before the presentation of a stimulus to facilitate later stimulus processing (Friston, 2005). As a consequence, one could expect the integration of pre-lexical familiarity of the learned pseudowords into the prediction process. If this were the case, we should have found the interaction of pre-lexical familiarity and context-based facilitation. Also, the predictive coding theory assumes a hierarchical architecture in which top-down besides within-level predictions facilitate processing (Friston, 2005). Once more, we could not find evidence for an architecture that implements top-down facilitation. However, we, on the other hand, provide evidence that a core mechanism of predictive coding (i.e., suppression of the informative part of expected sensory signals) is computationally implemented during visual word processing, to achieve context-based facilitation within, e.g., lexical processing levels.

Finally, one can speculate that specific lexical and pre-lexical information is transformed after completion of lexical access (i.e., after the N400 time window). The retrieved pre-lexical and lexical-semantic information might be used to predict the future stimuli already at the sensory level (e.g., Rao and Ballard, 1999). Note, predicting away information at the sensory level optimizes processing at later levels since, as prominently proposed in the predictive coding models, only the residual, i.e., unpredicted information, is processed at higher levels (e.g., Gagl et al., 2018). In line with this notion is that for visual and lexical processing a reduction of activation was found. Still, for the implementation of a sensory prediction as suggested here, the information has to be transformed, i.e., from lexical to visual information, and held active until the presentation of the next stimulus. At the prime, the late occipital context effect of higher activation for prime vs. target (i.e., C6, Fig. 5) might reflect the result of such a transformation process. We speculate that at this point in time top-down information might be used to prepare visual processing levels for the upcoming target presentation. When the subsequent target is in accordance with the prediction, based on predictive coding one can expect that the neuronal activation at the visual level is low. This expectation is met by the early context effect found in the occipital cortex (i.e., C1 cluster). Here, we expect that future research, e.g., using explicit connectivity investigations or specifically investigating the interval between prime and target, might allow specifying the information content integrated at the prime to facilitate processing of the target.

## Conclusion

In sum, our investigation of context-and familiarity-based facilitation of visual word recognition indicated within-level facilitation at visual and lexical processing levels. We found no support for hierarchical, top-down facilitation from a predictive (higher-level) context (e.g., word semantics) to lower levels of processing (visual, pre-lexical). At a mechanistic level, we could identify predictive coding as the most likely candidate for the implementation of facilitation processes (as compared to fatigue and sharpening). A novel approach of our study was the explicit manipulation of pre-lexical (i.e., orthographic and phonological) familiarity, via a pseudoword familiarization training procedure. We could not find support for context-based facilitation at the pre-lexical level but could identify a context-independent pre-lexical familiarity effect in the left angular/ supramarginal gyrus. Thus, we conclude that context-based facilitation relies on information about visual and lexical-semantic features of upcoming words. Interestingly, in natural reading, visual information is typically available through para-foveal pre-processing (Gagl et al., 2014; Schotter et al., 2012), while lexical-semantic information is available through previous text or sentence context (Hawelka et al., 2010; Kliegl et al., 2006). This analogy might indicate that context-based facilitation in reading mainly operates by visual and lexical representations. Investigating this dichotomy further in future studies might provide exciting avenues for refining the understanding of contextual influences on efficient word recognition during reading.

## Acknowledgements

The research leading to these results has received funding from the European Community’s Seventh Framework Programme (FP7/2013) under grant agreement n° 617891 awarded to C.J.F. and a Marie Curie fellowship under grant agreement n° 707932 awarded to B.G.. We thank Anne Hoffmann, Cordula Hunt, Jan Jürges, Rebekka Tenderra, and Stefanie Wu for their help with data acquisition as well as Klara Gregorova, Rebecca Anik Mayer, and Jona Sassenhagen for giving helpful comments on previous versions of the manuscript.

## Author Contributions

Experiment 1: C.J.F. and B.G. designed research; S.E. and B.G. performed research and analyzed data. Experiment 2: S.E., C.J.F., and B.G. designed research; S.E. and B.G. performed research and analyzed data. S.E., C.J.F., and B.G. wrote the paper. Authors report no conflict of interest.

## Extended Data for Figures

**Figure 3-1.**
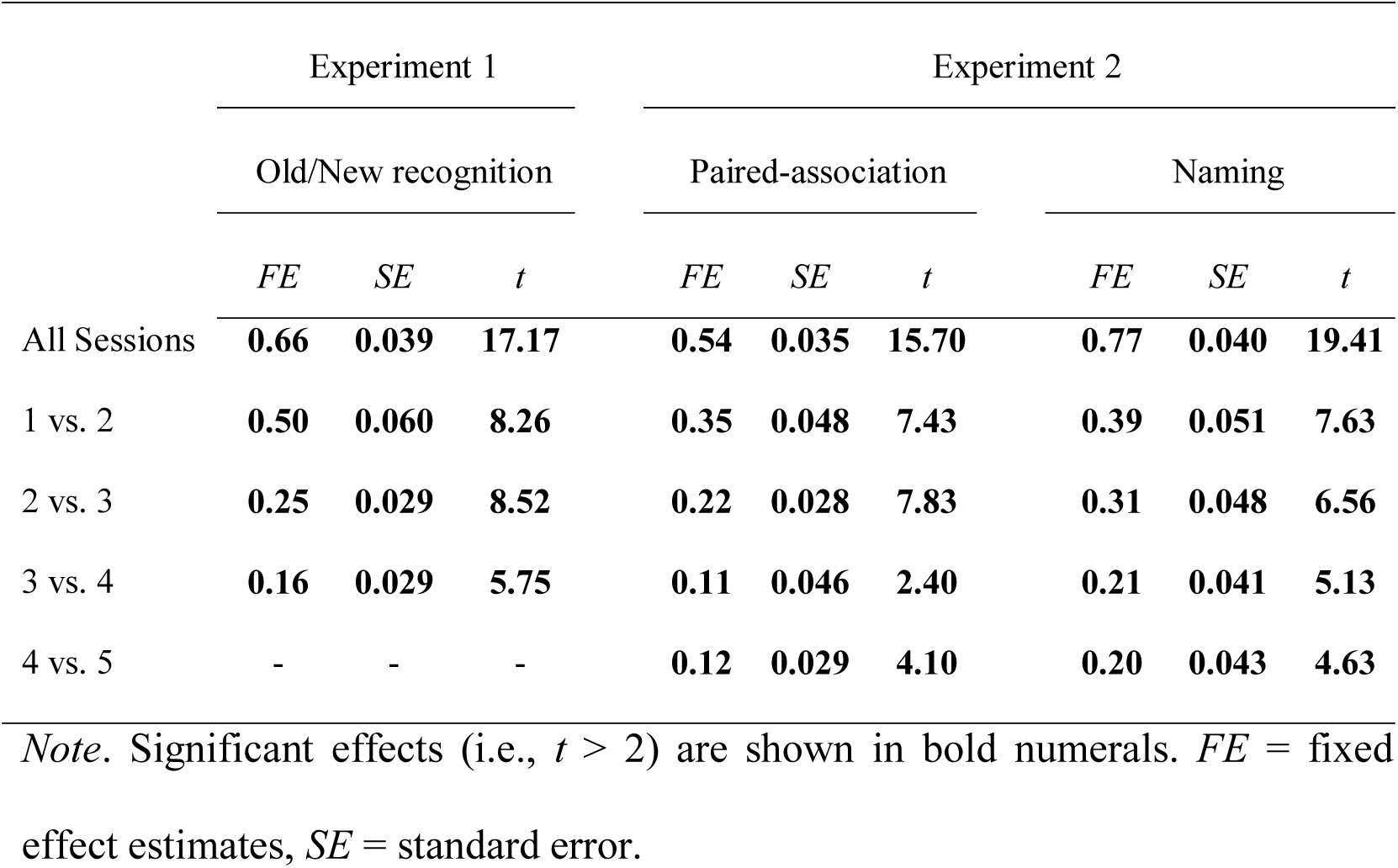
Results from the linear mixed model analyses of d’ sensitivity indices from the pseudoword familiarization sessions of both Experiment 1 and 2, including pairwise comparisons from session to session.

**Figure 4-1.**
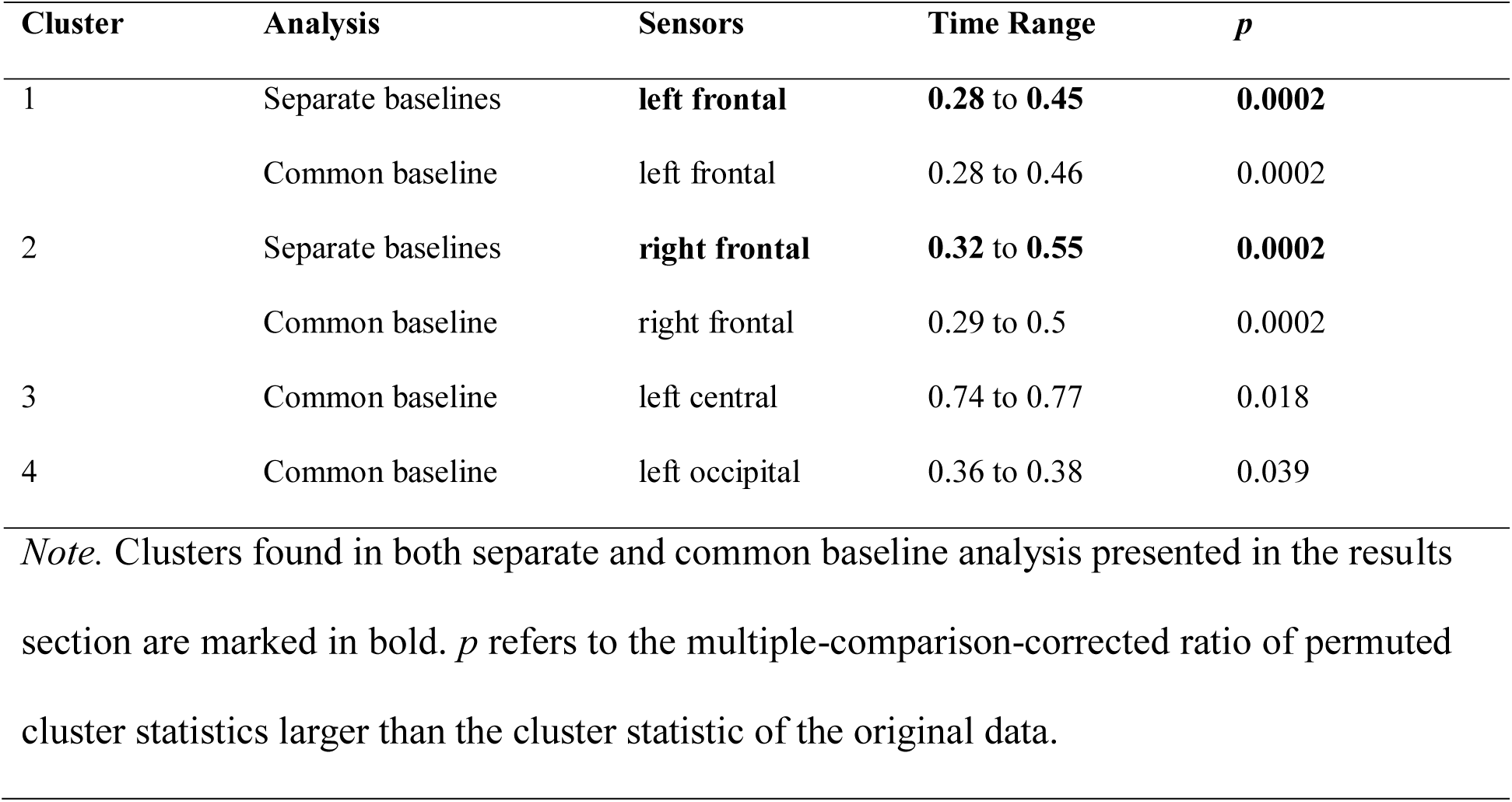
Overview of clusters demonstrating a prime/target by repetition congruency (repetition vs. non-repetition) interaction obtained with separate or common baselines for prime and target.

**Figure 5-1.**
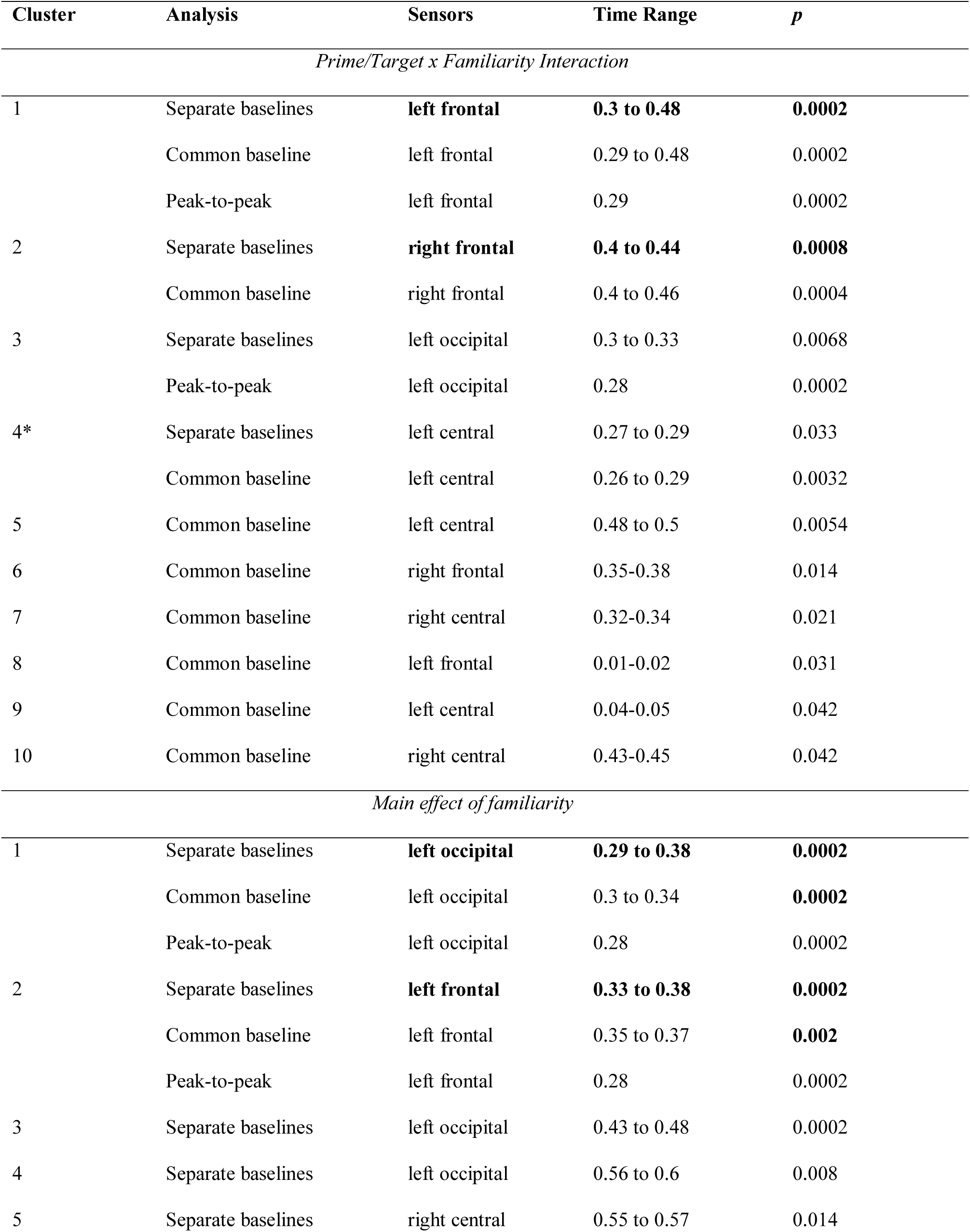
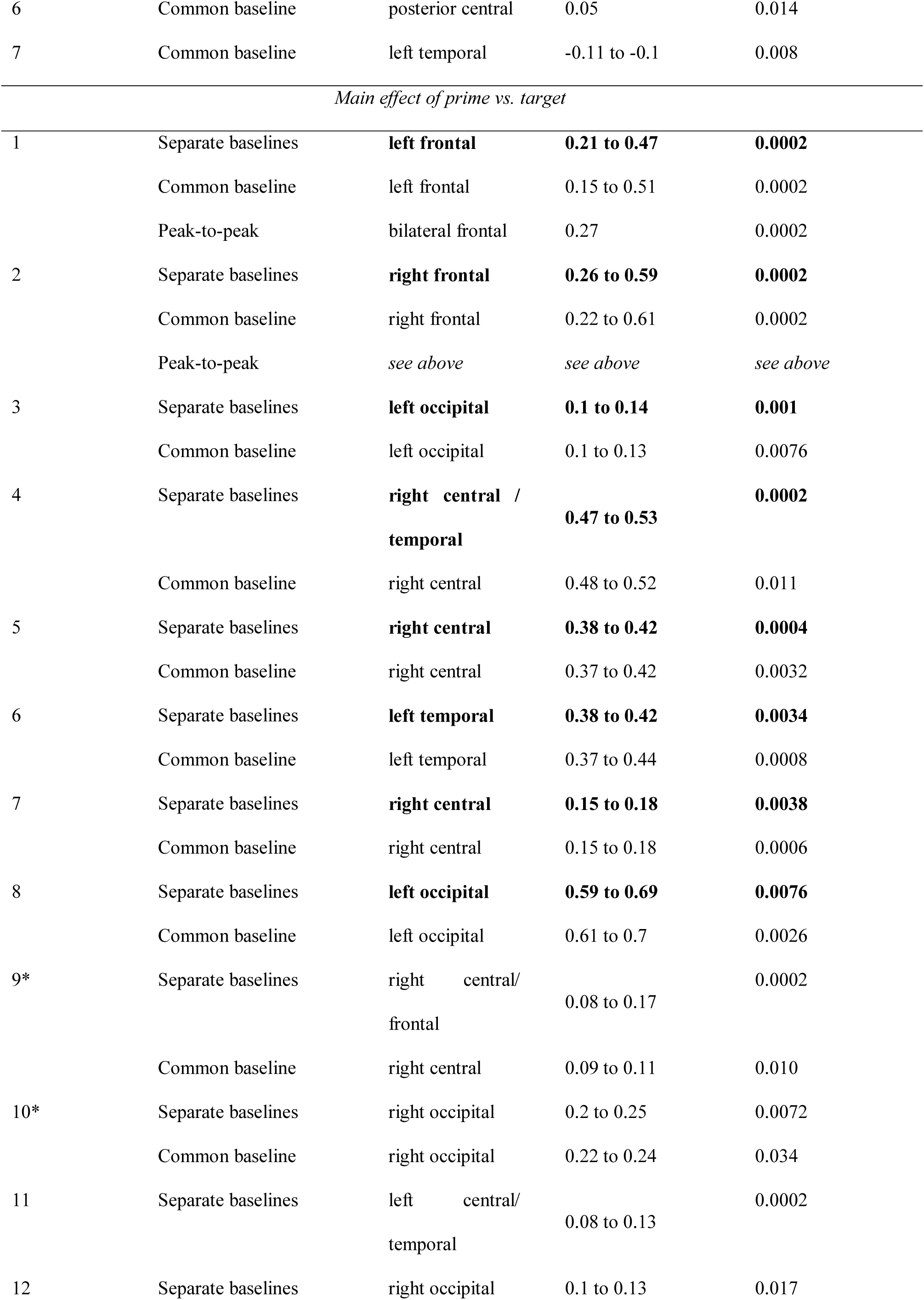
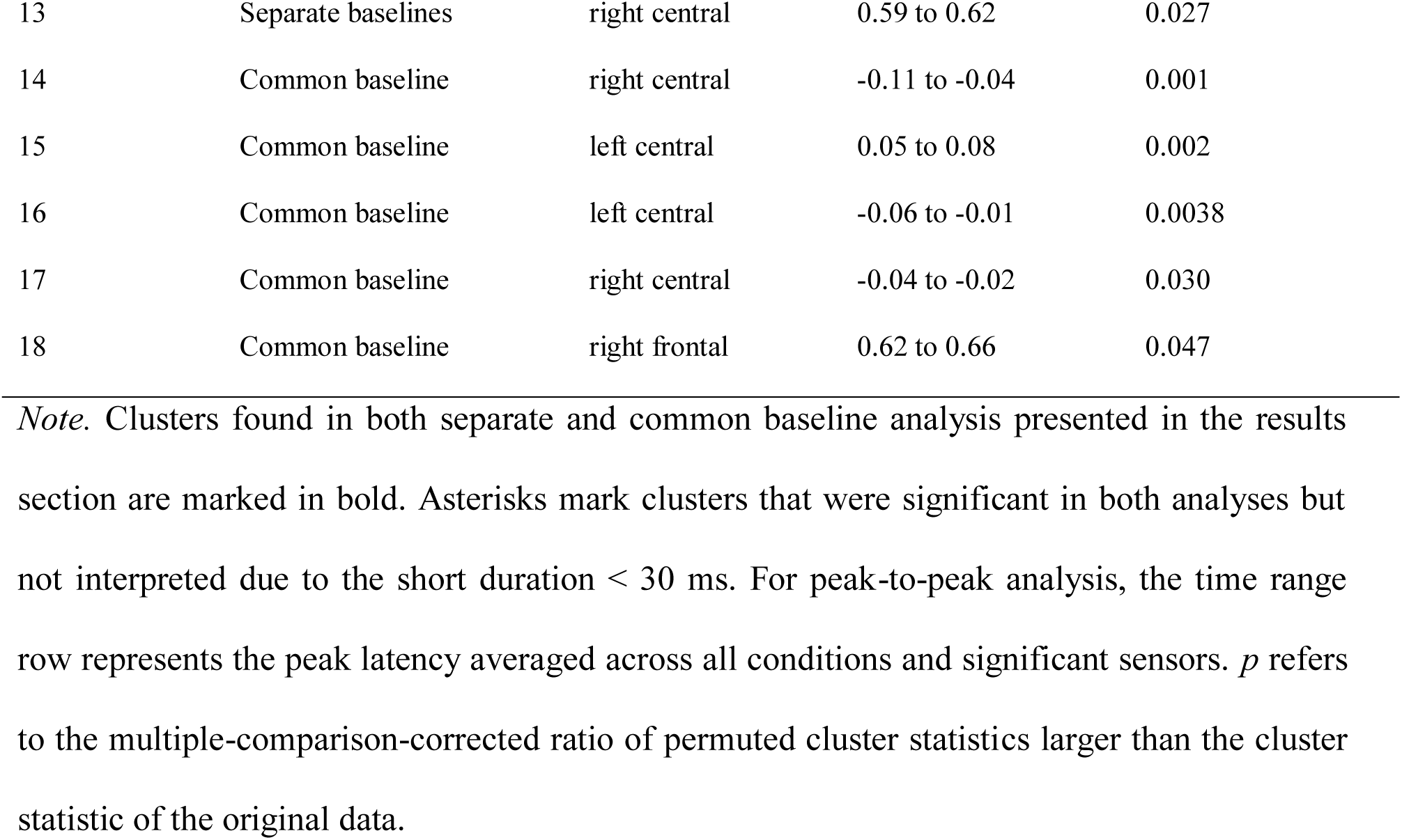
Overview of clusters form the analysis investigating the prime/target x familiarity interaction obtained with separate or common baselines for prime and target, as well as peak-to-peak analysis.

**Figure 5-2.**
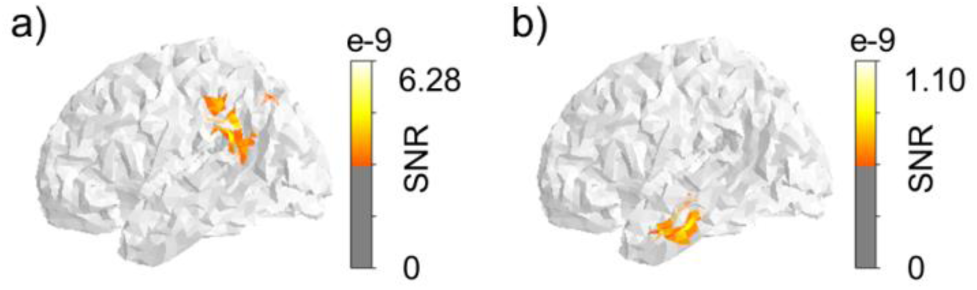
Source locations for a) the familiarity cluster F1 contrasting familiar pseudowords > words and b) the interaction cluster CxF contrasting prime vs. target by words vs. familiar pseudowords.

**Figure 5-3.**
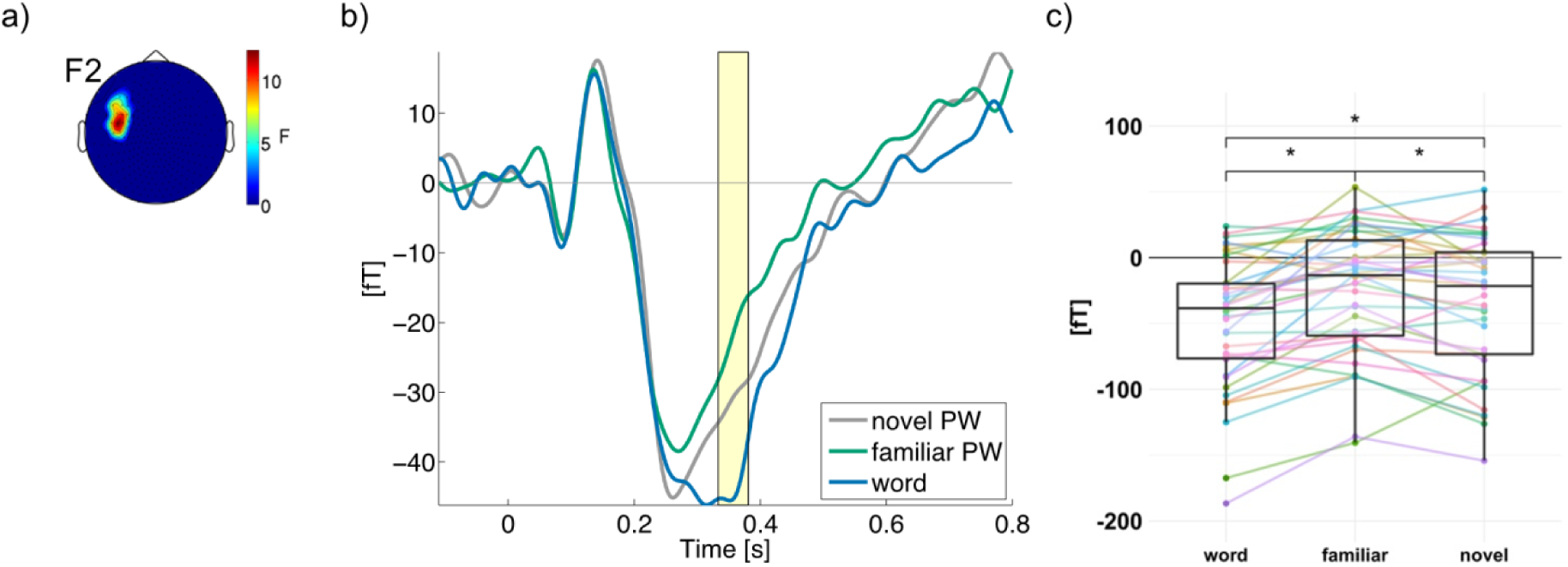
Main effect of lexical familiarity. a) Topographical map, b) ERF time course, and c) boxplot, averaged across prime and target.

**Figure 5-4.**
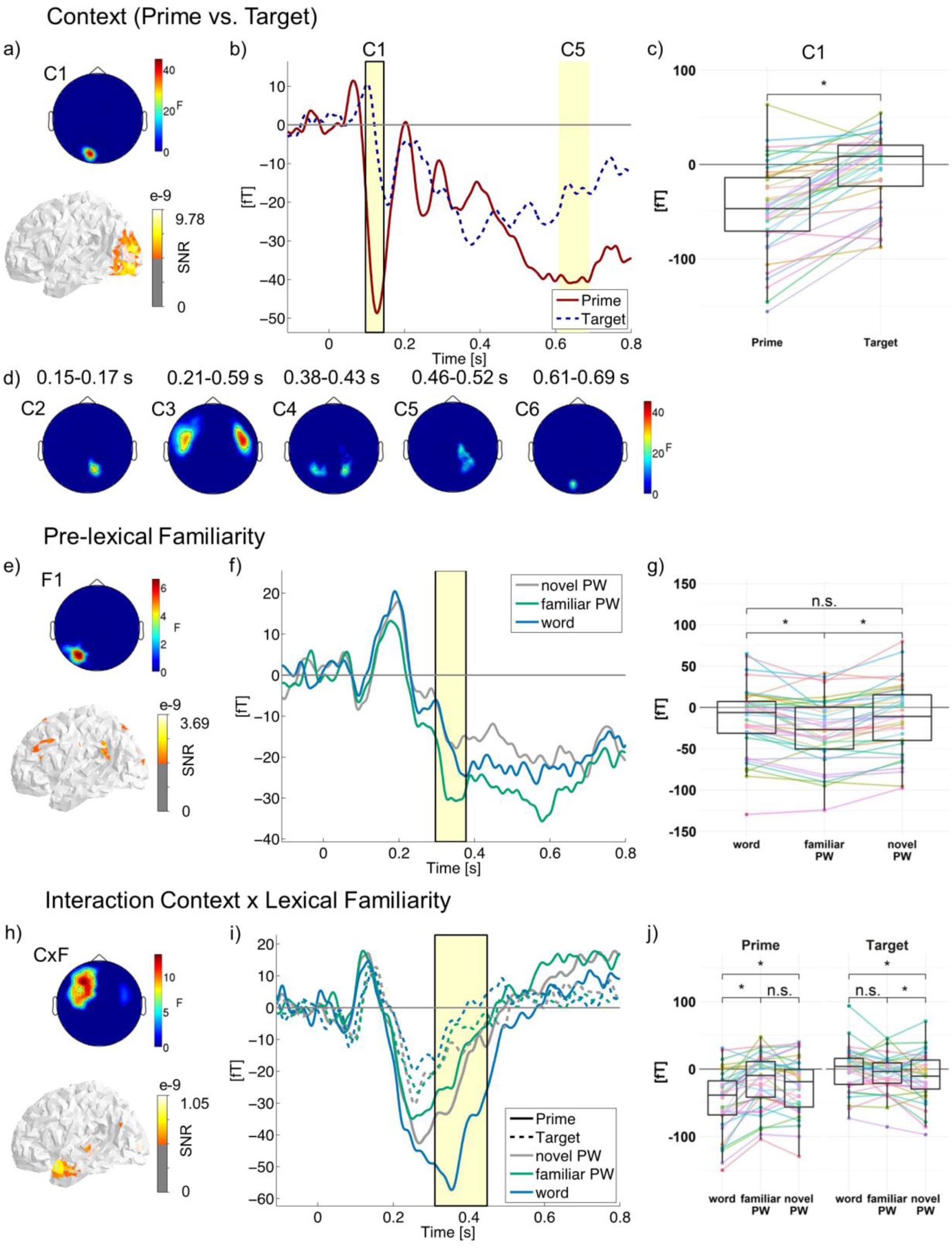
Same results as in Fig. 5 depicted for a low-pass filter of 40 instead of 20 Hz.

**Figure 5-5.**
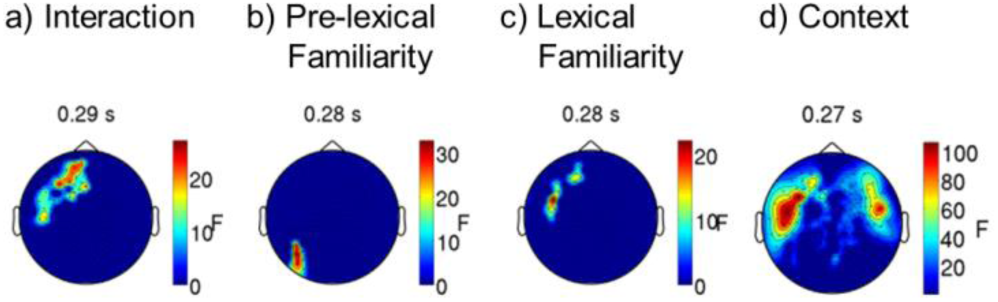
Significant clusters obtained in the peak-to-peak analysis for a) the interaction of context (prime vs. target) by lexical familiarity, b) main effects of pre-lexical familiarity, c) main effects of lexical familiarity and d) main effect of context (prime vs. target). Topographical maps represent *F*-values of significant sensors; note the different scales. Non-significant sensors are set to zero. Peak latencies averaged across significant sensors and all conditions are depicted above the topographical maps.

**Figure 6-1.**
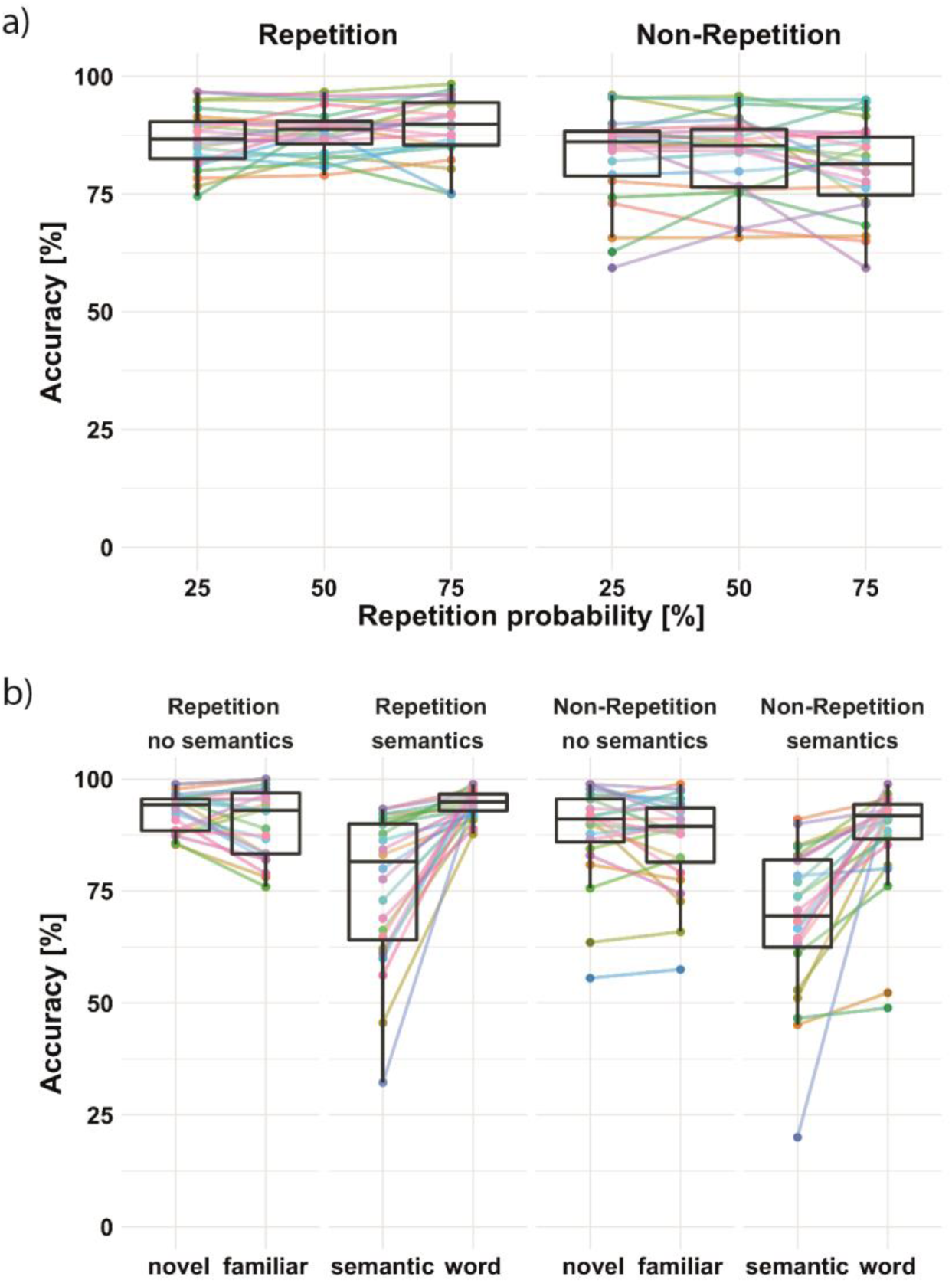
Accuracies in Experiment 2 (semantic association judgment task). a) Repetition probability effect for repetition (left) and non-repetition trials (right) averaged across familiarity conditions. b) Familiarity influence on context effects, i.e., repeated (left) vs. non-repeated targets (right). Effects are separated for familiarity conditions, with an additional separation for letter strings with and without semantic associations, averaged across repetition probabilities. Colored dots and lines represent individual participants.

## Extended Data for Tables

**Table 4-1.** Results from post hoc linear mixed model analyses on ERF values (in 10^-14^ Tesla) from sensor and time point of the strongest effect, separately for prime and target, for pre-lexical (F1) and lexical (F2) familiarity clusters represented in Fig. 5d-i and Extended Data Fig. 5-4, respectively.

|  | Prime |  |  | Target |  |  |
| --- | --- | --- | --- | --- | --- | --- |
|  | *FE* | *SE* | *t* | *FE* | *SE* | *t* |
|  | *F1* |  |  |  |  |  |
| Word vs. novel PW |  |  |  |  |  |  |
| Familiarity | -0.95 | 0.50 | 1.93 | **1.31** | **0.46** | **2.89** |
| OLD20 | -0.25 | 0.24 | 1.04 | -0.0059 | 0.22 | 0.026 |
| Number of syllables | -0.097 | 0.24 | 0.40 | -0.0046 | 0.22 | 0.021 |
| Words vs. familiar PW |  |  |  |  |  |  |
| Familiarity | **-2.59** | **0.48** | **5.44** | 0.052 | 0.46 | 0.11 |
| OLD20 | -0.057 | 0.24 | 0.24 | 0.023 | 0.23 | 0.10 |
| Number of syllables | -0.12 | 0.23 | 0.51 | -0.028 | 0.22 | 0.13 |
| Familiar vs. novel PW |  |  |  |  |  |  |
| Familiarity | **1.65** | **0.42** | **3.92** | **1.32** | **0.47** | **2.78** |
| OLD20 | -0.038 | 0.21 | 0.18 | -0.11 | 0.24 | 0.48 |
| Number of syllables | 0.35 | 0.21 | 1.67 | -0.019 | 0.24 | 0.081 |
| *F2* |  |  |  |  |  |  |
| Word vs. novel PW |  |  |  |  |  |  |
| Familiarity | **3.44** | **0.84** | **4.10** | -1.00 | 0.71 | 1.40 |
| OLD20 | -0.11 | 0.41 | 0.27 | -0.049 | 0.35 | 0.14 |
| Number of syllables | 0.12 | 0.42 | 0.30 | -0.092 | 0.35 | 0.26 |
| Words vs. familiar PW |  |  |  |  |  |  |
| Familiarity | **5.33** | **0.81** | **6.60** | -0.18 | 0.65 | 0.27 |
| OLD20 | 0.23 | 0.41 | 0.56 | -0.24 | 0.33 | 0.74 |
| Number of syllables | 0.30 | 0.39 | 0.77 | -0.30 | 0.31 | 0.94 |

|  | Familiar vs. novel PW |  |  |  |  |  |
| --- | --- | --- | --- | --- | --- | --- |
| Familiarity | **-1.66** | **0.67** | **2.46** | -0.81 | 0.60 | 1.35 |
| OLD20 | 0.035 | 0.34 | 0.10 | -0.37 | 0.30 | 1.26 |
| Number of syllables | 0.024 | 0.34 | 0.069 | -0.25 | 0.30 | 0.84 |

*Note.* Significant effects (i.e., $t > 2$ ) are shown in bold numerals. *FE* = fixed effect estimates, *SE* = standard error.

**Table 5-1.** Results from post hoc linear mixed model analyses on ERF values (in 10^-14^ Tesla) from sensor and time point of the strongest effect, separately for prime and target, for the prime/target x lexical familiarity interaction cluster represented in Fig. 5h-j.

|  | Prime |  |  | Target |  |  |
| --- | --- | --- | --- | --- | --- | --- |
|  | *FE* | *SE* | *t* | *FE* | *SE* | *t* |
| Word vs. novel PW |  |  |  |  |  |  |
| Familiarity | **2.15** | **0.47** | **4.54** | **-1.20** | **0.37** | **3.23** |
| OLD20 | -0.099 | 0.23 | 0.43 | 0.091 | 0.18 | 0.50 |
| Number of syllables | 0.11 | 0.23 | 0.46 | 0.026 | 0.18 | 0.14 |
| Words vs. familiar PW |  |  |  |  |  |  |
| Familiarity | **3.07** | **0.42** | **7.39** | -0.65 | 0.37 | 1.76 |
| OLD20 | 0.19 | 0.2101 | 0.91 | -0.17 | 0.19 | 0.90 |
| Number of syllables | 0.14 | 0.20 | 0.71 | 0.045 | 0.18 | 0.25 |
| Familiar vs. novel PW |  |  |  |  |  |  |
| Familiarity | -0.70 | 0.40 | 1.75 | -0.66 | 0.33 | 1.98 |
| OLD20 | -0.023 | 0.20 | 0.11 | -0.061 | 0.17 | 0.37 |
| Number of syllables | 0.038 | 0.20 | 0.19 | -0.15 | -0.17 | 0.91 |

*Note.* Significant effects (i.e., $t > 2$ ) are shown in bold numerals. *FE* = fixed effect estimates, *SE* = standard error.

**Table 10-1.** Results from the linear mixed model analyses investigating repetition congruency (repetition vs. non-repetition), probability, familiarity, and semantics in accuracies during repetition priming (Experiment 2).

|  | *FE* | *SE* | *z* | *p* |
| --- | --- | --- | --- | --- |
| Repetition congruency | **0.29** | **0.027** | **10.78** | **< 2e-16** |
| Probability | 0.023 | 0.028 | 0.82 | 0.41 |
| Pre-lexical familiarity | **-0.44** | **0.027** | **16.08** | **< 2e-16** |
| Lexical familiarity | **-0.27** | **0.027** | **-9.8** | **< 2e-16** |
| Repetition congruency x Probability | **0.10** | **0.028** | **3.58** | **0.00034** |
| Repetition congruency x Pre-lexical familiarity | **-0.09** | **0.027** | **3.31** | **0.00093** |
| Repetition congruency x Lexical familiarity | 0.027 | 0.027 | 1.00 | 0.32 |
| Probability x Pre-lexical familiarity | -0.021 | 0.028 | -0.76 | 0.44 |
| Probability x Lexical familiarity | **0.064** | **0.028** | **2.32** | **0.020** |
| Pre-lexical x Lexical familiarity | **-0.28** | **0.027** | **10.38** | **< 2e-16** |
| Repetition congruency x Probability x Pre-lexical familiarity | -0.037 | 0.028 | 1.35 | 0.18 |
| Repetition congruency x Probability x Lexical familiarity | **0.075** | **0.028** | **2.72** | **0.0065** |
| Repetition congruency x Pre-lexical x Lexical familiarity | -0.039 | 0.027 | 1.44 | 0.15 |
| Probability x Pre-lexical x Lexical familiarity | -0.015 | 0.028 | 0.54 | 0.59 |
| OLD20 | **0.071** | **0.023** | **3.06** | **0.0022** |
| Number of syllables | 0.045 | 0.024 | 1.84 | 0.065 |

*Note.* Significant effects (i.e., $p < 0.05$ ) are shown in bold numerals. *FE* = fixed effects, *SE* = standard error.

**Table 10-2.**
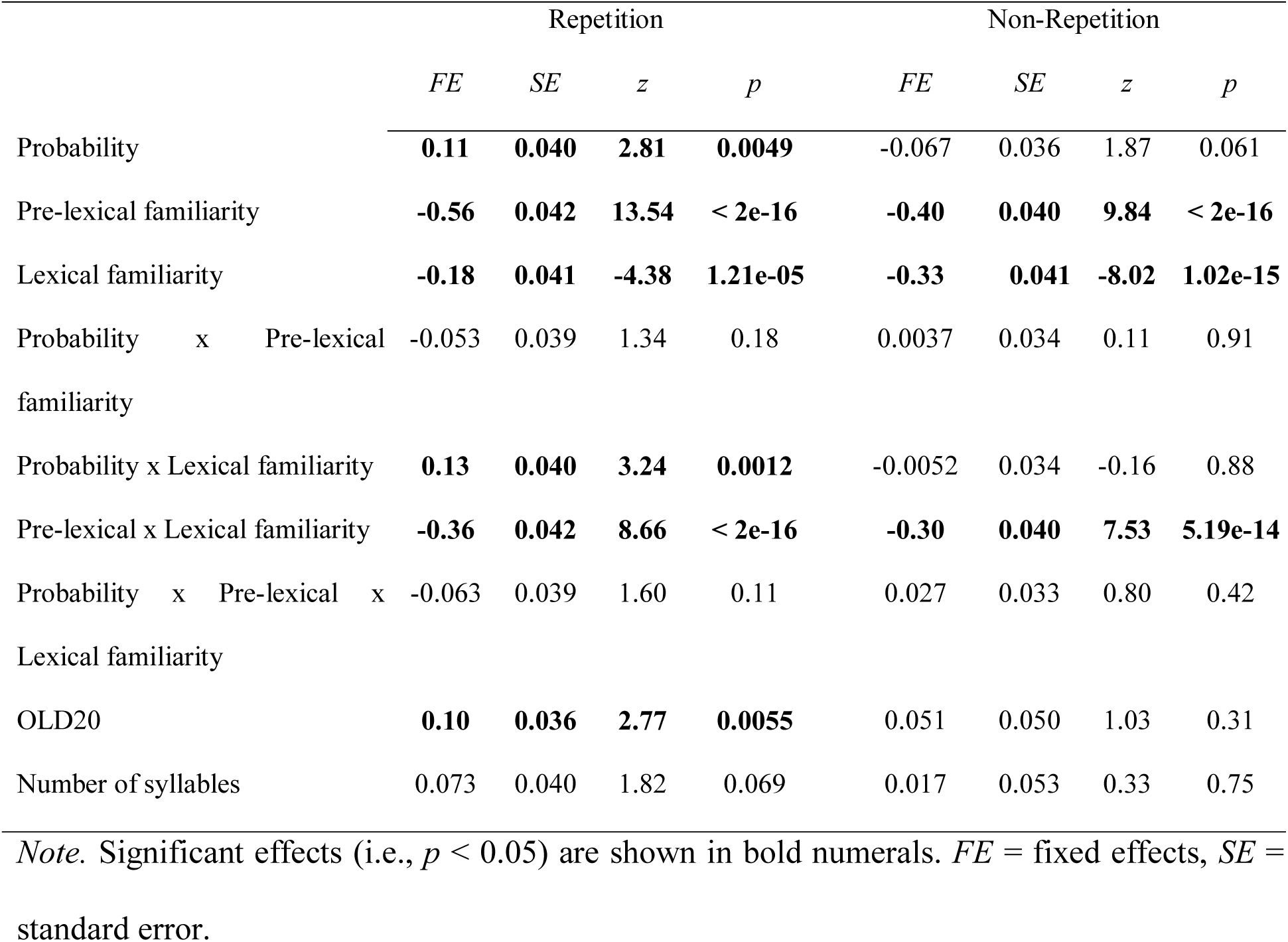
Results from the linear mixed model analyses on accuracies during repetition priming (Experiment 2), separately for repetition and non-repetition trials.

